# PHOSPHATIDYLSERINE EXPOSURE AND EXTRACELLULAR ANNEXIN A5 WEAKEN THE ACTIN CORTEX IN OSTEOCLAST FUSION

**DOI:** 10.1101/2025.07.16.663971

**Authors:** Evgenia Leikina, Andrey K. Tsaturyan, Kamran Melikov, Jarred M. Whitlock, Jared Cunanan, Morgan Roegner, Griffin Katz, Michael M. Kozlov, Leonid V. Chernomordik

**Author notes:** Correspondence: Drs. Leonid V Chernomordik and Michael M. Kozlov.

## Abstract

Diverse cell-cell fusions involve intracellular Ca^2+^ signaling, non-apoptotic exposure of phosphatidylserine (PS) at the surface of fusion-committed cells and binding of extracellular Annexin A5 (Anx A5). Here we focus on the cell fusion stage of formation of bone-resorbing multinucleated osteoclasts and report that each of the listed hallmarks of cell fusion represents a step in a novel bidirectional signaling pathway. A rise in intracellular Ca²⁺ activates a lipid scramblase that translocates PS from the inner to the outer leaflet of the plasma membrane. This redistribution is enhanced by extracellular Anx A5 binding to cell surface PS. Depletion of PS in the inner leaflet weakens actin cortex-plasma membrane attachment mediated by ezrin/radixin/moesin (ERM) proteins, as evidenced by the preferential localization of cortex detachment areas within PS-enriched regions at the surface of the cells. Weakening of the cortex-membrane connection by Anx A5 or by adding an inhibitor of the ERM proteins promotes osteoclast fusion. We propose that this pathway facilitates osteoclast fusion and other cell-cell fusions by promoting membrane deformations required for formation of prefusion membrane contacts. Additionally, the elevated tension in the cortex detachment region of the membrane, suggested by our theoretical analysis, promotes fusion pore expansion.

## INTRODUCTION

Cell-cell fusion plays a critical role in fertilization, the lifelong remodeling of bones and skeletal muscles, the formation of the placental syncytiotrophoblast, and other physiological processes ^1^. Different fusion processes involve distinct protein regulators, including Myomaker and Myomerger (also referred to as Myomixer and Minion) in myoblast fusion ^2^, Syncytins in trophoblast fusion ^3,4^, and Fusexins in C. *elegans*^1^. In contrast to these fusion-process-specific proteins, which are required and sufficient for fusion and are thought to be directly involved in the remodeling of membrane bilayers, some mechanistic motifs appear to be strikingly conserved across diverse cell-cell fusion processes. Myoblast, trophoblast, and osteoclast fusion rely on calcium signaling ^5–7^ and on lipid scramblases, proteins that mediate the nonselective, bidirectional movement of lipids between the inner (cytofacial) and outer leaflets of the plasma membrane (PM) ^8–11^. Diverse cell fusions are found to depend on non-apoptotic cell surface exposure of an anionic lipid phosphatidylserine (PS), normally almost exclusively retained in the inner leaflet of the PM (reviewed in ^12^). These fusions also rely on extracellular PS-binding proteins and, specifically, Annexin A5 (Anx A5) (reviewed in ^12^ and ^13,14^). Another mechanistic motif shared across diverse fusion processes—including myoblast fusion ^15,16^, trophoblast fusion ^17^, macrophage fusion during osteoclast formation (reviewed in ^18^ and ^19,20^), and in multinucleated giant cells ^21^ —is the formation of outward-going PM protrusions ^22,23^. The best characterized example of such membrane structures is invadosomes in myoblasts ^4,21^.

Local deformations of the PM in protrusions, including thin (60–200 nm diameter) finger-like filopodia ^24,25^ and blister-like spherical membrane blebs, are regulated by actin cytoskeleton remodeling and/or positive intracellular pressure generated by actomyosin contractility and directional water flux across the PM ^26^. Initiation of PM deformations depends on the local and transient detachment of a patch of membrane from the actin cortex (AC), membrane-linked actin-rich protein network, and/or dissociation of this network ^27,28^. The thickness, density, and dynamics of the AC, as well as its subcellular distribution, vary between cell types, depend on the physiological state of the cell, and are regulated by numerous actin-binding proteins^29^. AC is attached to the inner leaflet of the PM via membrane-anchoring proteins of the ezrin/radixin/moesin (ERM) family ^30^. ERM binding to the PM and organization of the actomyosin network depend on the membrane lipid composition and are promoted by PS and another anionic lipid phosphatidylinositol 4,5-bisphosphate (PI(4,5)P2) ^31–34^. Like PS, PI(4,5)P2 is highly enriched in the inner leaflet of PM ^35,36^, but in some physiological processes it is also found at the cell surface ^37,38^. Local weakening of the AC-PM attachment, via mechanisms such as reduction of the negative surface charge of the inner leaflet, allows intracellular pressure to locally bulge the PM ^27,39–42^. Several studies have suggested that cell fusion depends on weakening of the PM-associated AC ^20,43,44^. However, the specific mechanisms linking cell fusion to AC weakening or detachment and to the formation and properties of outward PM protrusions, remain to be clarified.

Why do diverse cell fusion processes depend on lipid scramblase activity and on the appearance of PS- and Anx A5 at the cell surface? In this study we address this question for osteoclast fusion, a critical stage in the formation of bone-resorbing multinucleated osteoclasts ^45^. Each fusion event increases the bone-resorbing activity of osteoclasts, which balances the bone-forming activity of osteoblasts in the tightly regulated, lifelong remodeling of the skeleton. The mechanisms of osteoclast fusion and its importance in bone maintenance and disease have been explored in many recent studies (reviewed in ^46–49^). Here, we report that activation of lipid scrambling reduces PS and PI(4,5)P₂ levels in the inner leaflet of the PM, leading to local AC–PM detachment. Binding of extracellular Anx A5 to cell-surface PS enhances PS depletion from the inner leaflet and thus further weakens the AC-PM connection. We propose that detachment of the PM from the AC facilitates tight contacts between lipid bilayers of the two PMs and promotes their fusion. Our findings may help explain why osteoclast fusion, and possibly other cell–cell fusion processes, depend on PS externalization, extracellular Anx A5 and membrane protrusions.

## RESULTS

### Cell-surface Anx A5 controls osteoclast fusion

Osteoclast differentiation is accompanied by a transient loss of lipid asymmetry in the PM, manifested as the appearance of PS at the surface of osteoclasts ^8^. In turn, this PS exposure increases the presence of endogenous extracellular Anx A5 at the cell surface ^8^. Reducing the expression of endogenous Anx A5 and suppressing its function at the cell surface using antibodies to Anx A5 and peptide inhibitors reduces osteoclast fusion ^8,14^. To test whether the amounts of cell-surface-associated Anx A5 limits the efficiency of osteoclast fusion, we examined the effects of applying recombinant Anx A5 (rAnx A5). First, we verified that this application increases the total amount of Anx A5 at the cell surface, as evidenced by an increase in antibodies to Anx A5 staining (Fig. 1A,B). This finding suggests that endogenous extracellular Anx A5 is insufficient to saturate all surface-exposed PS, leaving some externalized PS accessible for rAnx A5 binding. We then explored the effects of rAnx A5 application specifically on the cell fusion stage of formation of multinucleated osteoclasts, uncoupled from the pre-fusion stages of osteoclastogenic differentiation, using cell fusion synchronization approach ^8,14,20,50^. We accumulated ready-to-fuse osteoclasts in the presence of a fusion inhibitor lysophosphatidylcholine (LPC), then washed out the inhibitor to allow fusion to proceed. Application of rAnx A5 at the time of LPC removal resulted in a robust promotion of synchronized osteoclast fusion (Fig. 1C,D). These data indicate that surface-bound Anx A5 in fusion-committed osteoclasts positively regulates the efficiency of fusion.

**Fig. 1:**
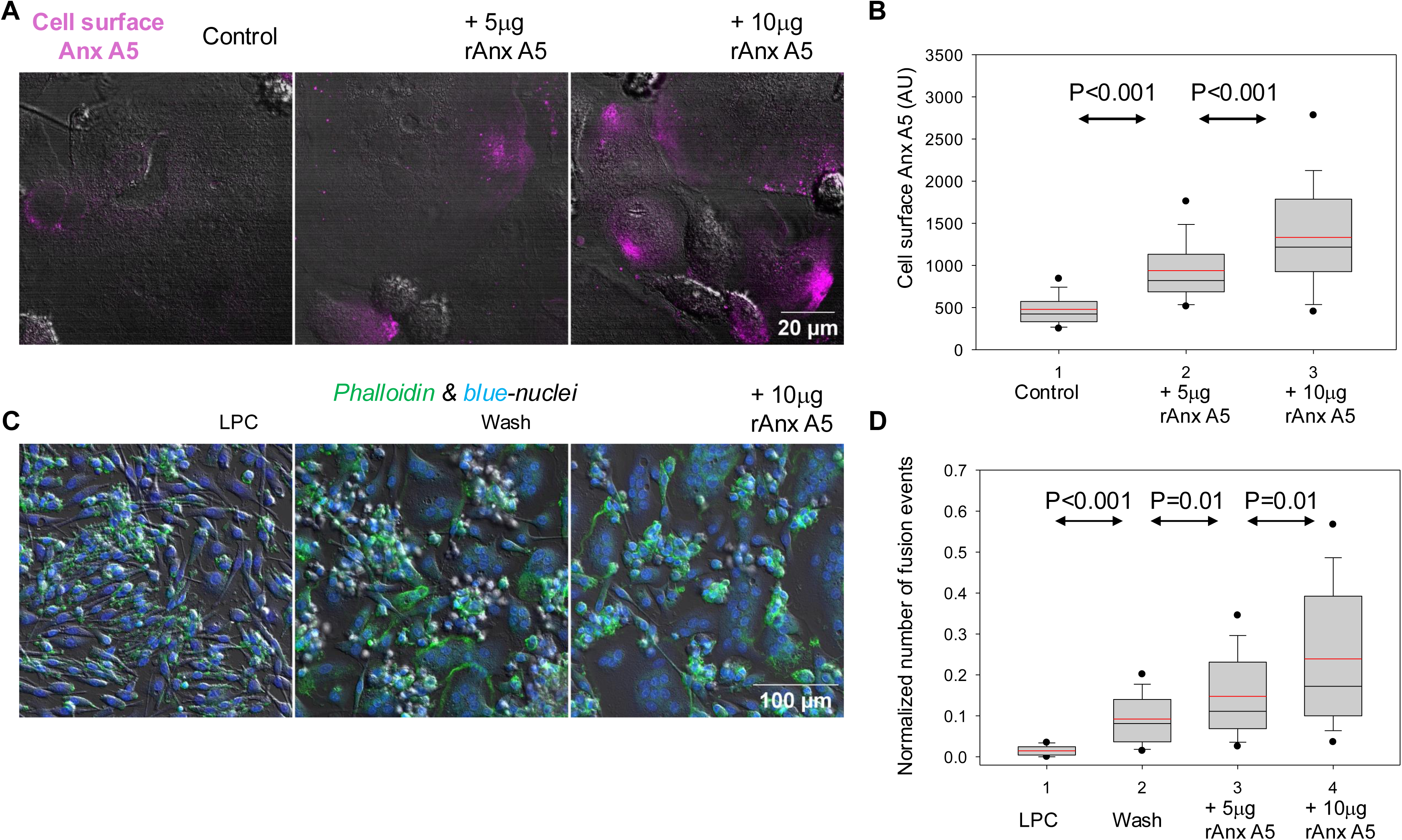
Cell surface Anx A5 controls the efficiency of osteoclast fusion. **A,B**. Representative fluorescence microscopy images (**A**) and quantification (**B**) demonstrate increased surface levels of Anx A5 on differentiating, non-permeabilized human osteoclasts four days after RANKL application, following the addition of unlabeled recombinant Anx A5 (rAnx A5). 10 min after rAnx A5 application, the cells were washed three times with complete medium and fixed. To detect both endogenous and exogenously applied Anx A5, cells were stained with antibodies to Anx A5 followed by fluorescently labeled secondary antibodies (shown in magenta). The amount of cell-surface bound Anx A5 was quantified by measuring total cell-associated fluorescence. **C, D**. Application of unlabeled rAnx A5 promotes osteoclast fusion decoupled from pre-fusion differentiation stages. Fusion was synchronized with the hemifusion inhibitor lysophosphatidylcholine (LPC), applied for 16 hours starting three days after RANKL treatment. rAnx A5 was applied at the time of LPC removal, when the cells were allowed to fuse. Representative fluorescence microscopy images (**C**) and quantification (**D**) of the numbers of cell fusion events at 2 hours after LPC removal in the presence of 0 (“Wash”), 5 or 10 μg/ml rAnx A5. “LPC” – fusion observed without removal of LPC. **B, D**. Data in **B** and data in **D** are pooled each from 3 independent experiments (>3,000 cells for each condition; >30 fields for **B** and **D**, respectively). Data are presented as box plots with center line showing the median, red line showing the mean, and box limits indicating the 5^th^ and 95^th^ percentiles. Statistical significance was assessed via two-tailed t-test.

### Lipid scrambling reduces PS and PI(4,5)P2 concentrations in the inner leaflet

Activated lipid scramblase facilitates the redistribution of anionic lipids —normally enriched in the inner leaflet of the PM—to the outer leaflet. To examine whether this lipid redistribution is accompanied by a decrease in PS content in the inner PM leaflet, we used HEK293 cells engineered to express TMEM16F scramblase after doxycycline induction ^51^. We activated scramblase activity, and thus PS exposure at the surface of doxycycline-induced cells (TMEM16F-HEK293), by stimulating intracellular Ca²⁺ elevation with the ionophore ionomycin. To analyze intracellular PS distribution, we transfected TMEM16F-HEK293 cells with a GFP-tagged intracellular PS probe (LactC2), which binds PS independently of Ca²⁺. As in other mammalian cells ^52^, TMEM16F-HEK293 cells showed characteristic PM-enriched localization of the PS probe under control conditions (Fig. 2A, left image). In contrast, following TMEM16F activation via Ca^2+^ addition, scrambling cells, identified by surface staining with rAnx A5, exhibited mostly cytoplasmic localization of the PS probe (Fig. 2A, right). In similar experiments with TMEM16F-HEK293 cells expressing a CFP-tagged PI(4,5)P2 probe, activation of lipid scrambling also reduced PI(4,5)P2 content in the inner leaflet of the PM (Fig. 2B). This pronounced shift in the localization of PS and PI(4,5)P₂ probes from inner leaflet-associated to cytoplasm-associated was confirmed by quantification (Fig. 2C). Even after doxycycline and ionomycin/Ca^2+^ treatment, some cells showed no cell surface labeling with rAnx A5 (red) (Fig. 2D,E), suggesting that TMEM16F in these cells was either not induced, not delivered to the PM or not activated. As seen in Fig. 2D, E, scrambling cells and non-scrambling cells within the same field of view exhibit different intracellular distributions of PS and PI(4,5)P2, confirming the scrambling-dependent loss of these anionic lipids in the inner PM leaflet.

**Fig. 2:**
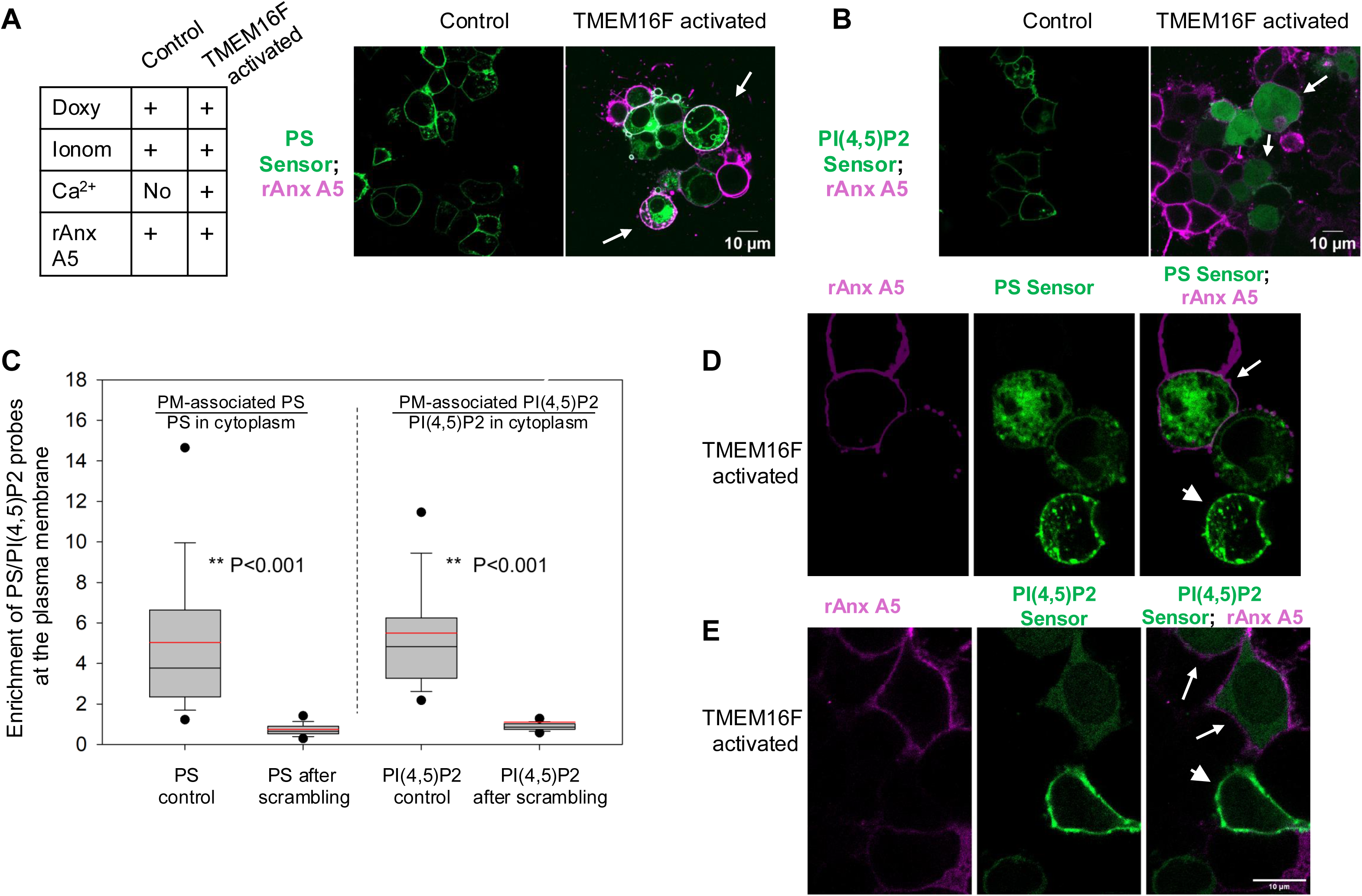
Lipid scrambling decreases PS and PI(4,5)P2 concentrations in the inner leaflet. **A, B.** Representative fluorescence microscopy images of TMEM16F-HEK293 cells expressing intracellular PS (**A**) or PI(4,5)P2 (**B**) probes. In both **A** and **B** images on the left show the characteristic PM labeling for the cells treated with doxycycline and then ionomycin and fluorescent rAnx A5 (magenta) applied in the absence of extracellular Ca^2+^ (Control). Images on the right show the cells with doxycycline induced TMEM16F activated by 5 min/37°C incubation with ionomycin, rAnx A5 and Ca^2+^. Images were taken 10 min after activation. The scrambling cells identified by the surface staining with rAnx A5 (magenta, marked by arrows) show cytoplasm-staining by PS- (**A**) and PI(4,5)P2- (**B**) probes. **C.** Quantification of the scrambling-induced shift from PM- to cytoplasmic-labeling with PS- and PI(4,5)P2-probes in the experiments such as the ones shown in **A** and **B**. Data (ratios of mean per pixel fluorescence in PM-proximal and cytoplasmic regions) for more than 45 cells for each condition pooled from 3 independent experiments and presented as box plots with center line showing the median, red line showing the mean, and box limits indicating the 5^th^ and 95^th^ percentiles. Statistical significance was assessed via two-tailed t-test. **D, E**. Comparison of intracellular distributions of PS- and PI(4,5)P2-probes in TMEM16F-HEK293 cells treated with doxycycline, ionomycin, Ca^2+^ and rAnx A5, for scrambling cells (rAnx A5 stained, marked by arrows, probes are mostly in cytoplasm) vs. not scrambling cells (no rAnx A5 staining, marked by arrowheads, probes are enriched at the inner leaflet of PM).

These findings indicate that activation of lipid scrambling decreases PS -and PI(4,5)P2-content in the inner leaflet of the PM, and that this decrease is not rapidly compensated by lipid biosynthesis or transport from the endoplasmic reticulum.

### Lipid scrambling and Anx A5 weaken the actin cortex-plasma membrane attachment

Given the dependence of ERM–PM binding on anionic lipids ^31–34^, a decrease in PS and PI(4,5)P2 content in the inner leaflet is expected to weaken the AC-PM attachment. Indeed, in experiments where we first labeled endogenous Anx A5 on the surface of non-permeabilized differentiating osteoclasts with antibodies to Anx A5, then permeabilized the cells and applied phalloidin to assess AC density, we observed diminished phalloidin labeling beneath the Anx A5-enriched PM patches (Fig. 3A). To evaluate the effects of PS exposure and extracellular Anx A5 on the AC in live cells, we visualized cell surface PS using rAnx A5 and labeled F-actin using one of two cell-permeable actin filament probes: CellMask Actin tracking stain (Fig. 3B) or SiR-Actin (Fig. 4A). Both probes have an actin-targeting structure that resembles jasplakinolide, but SiR-Actin binding to actin filaments is reversible. We observed scrambling- and Anx A5-dependent AC detachment (discussed below) with both probes, but quantified this effect only in experiments using CellMask Actin probe.

**Fig. 3:**
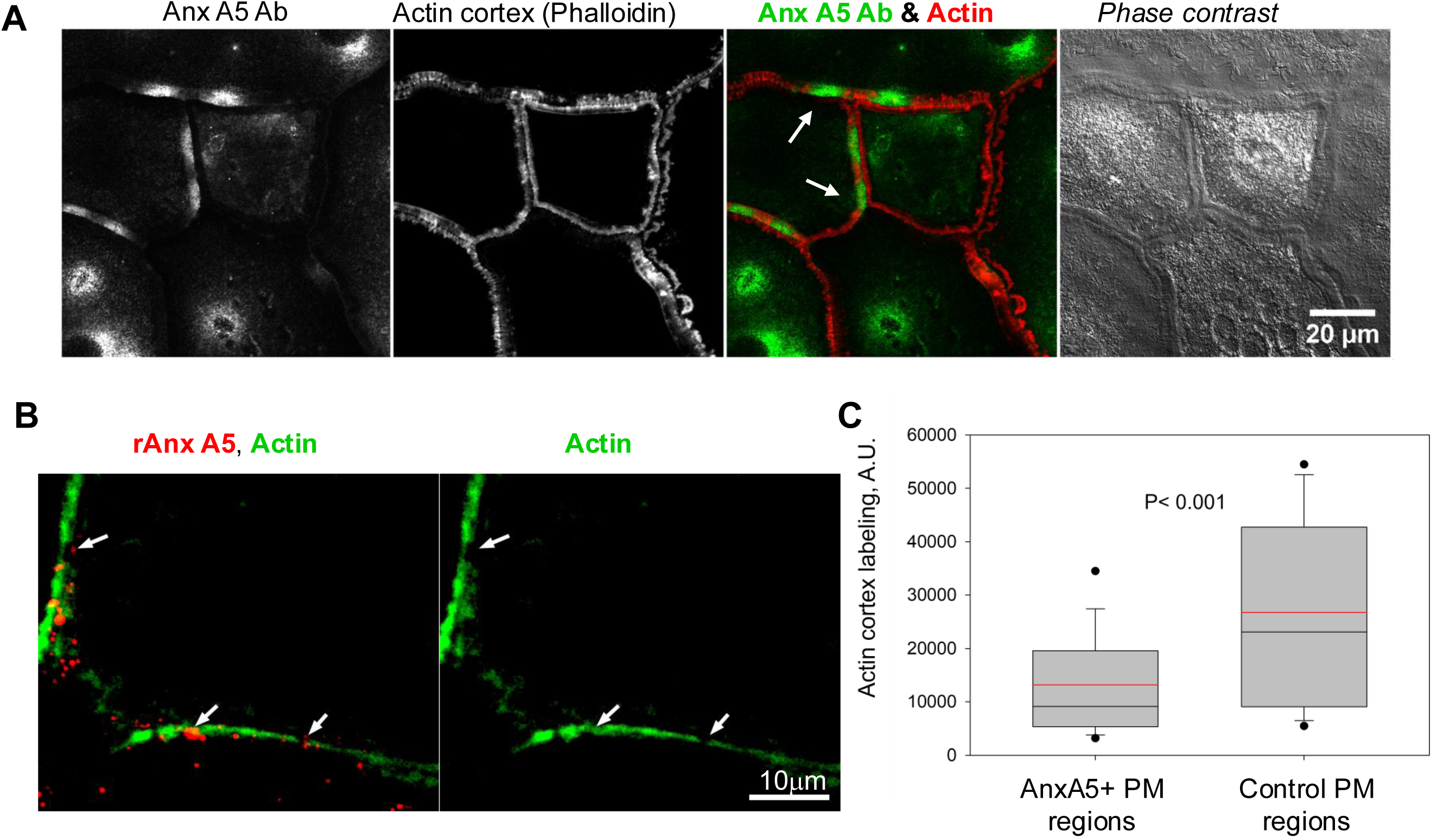
Lipid scrambling and extracellular Anx A5 weaken the actin cortex-plasma membrane attachment in differentiating osteoclasts. **A.** Representative fluorescence microscopy and phase contrast images of fixed osteoclasts at 4 days post-RANKL application stained with antibodies to Anx A5 to detect cell surface Anx A5 and then permeabilized and stained with phalloidin to detect filamentous actin. Arrows point to the PM regions with increased cell surface staining with antibodies to Anx A5 (green) and weaker actin filament staining (red). **B**. rAnx A5 (red)-enriched regions of PM of live differentiating osteoclasts 4 days post RANKL application (marked by arrows) show weaker labeling with Cellmask Green Actin (green). Images were taken 10 min after applying rAnx A5 at room temperature. **C**. Analysis of the experiments such as the one shown in **B**. Data (mean per pixel fluorescence of Cellmask Green Actin-labeled regions with clearly identifiable AC, labeled or not labeled by fluorescent rAnx A5) for more than 1700 measurements for each condition in 40 fields of views pooled from 7 independent experiments and presented as box plots with center line showing the median, red line showing the mean, and box limits indicating the 5^th^ and 95^th^ percentiles. Statistical significance was assessed via two-tailed t-test.

**Fig. 4:**
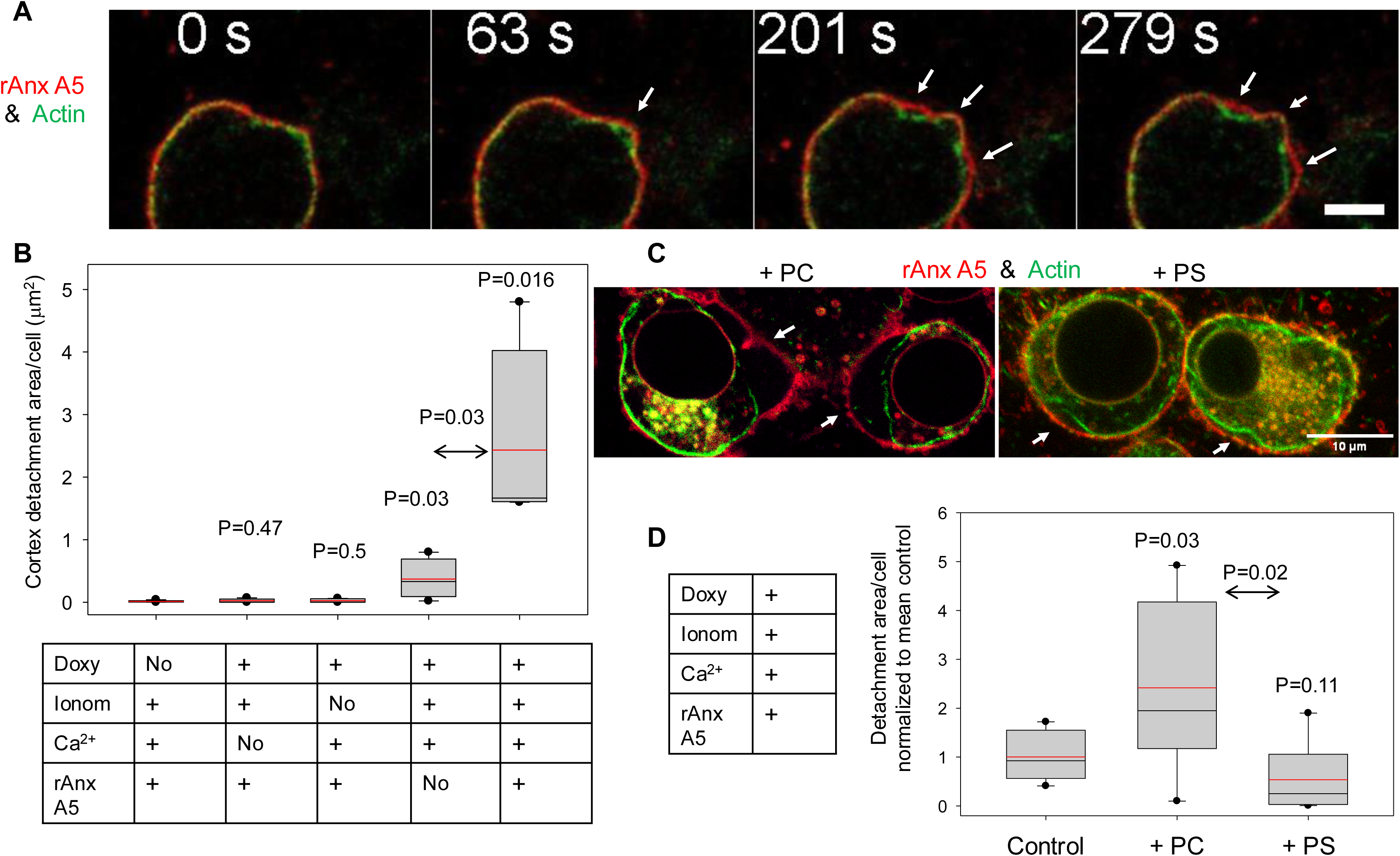
Lipid scrambling and Anx A5 binding to plasma membrane of TMEM16F-HEK cells promote actin cortex detachments. **A**. Representative fluorescence microscopy images (frames from the Supplemental Movie 1) of live TMEM16F-HEK cells induced to express TMEM16F and stained with cell-permeable F-actin probe SiR-Actin (green) and rAnx A5 (red) after ionomycin, rAnxA5 and Ca^2+^ application. The cells develop local cortex detachment areas (marked by arrows). t= 0 marks the application of rAnx A5. The scale bar is 5 μm. **B**. Quantification of cortex detachment area per cell for the experiments shown in Fig. S1) with the cells labeled with F-Actin probe Cellmask Green Actin and membrane probe FM 4-64 with activated TMEM16F scramblase with and without rAnx A5. In control experiments we either did not induce TMEM16F or not activated it by omitting ionomycin or Ca^2+^ application. Images were taken 10 min after Ca^2+^ and/or rAnx A5 application. Average cortex detachment area per cell was measured as described in the Methods for more than >1,200 cells for each condition. The data were pooled from 4 independent experiments and presented as box plots with center lines showing the medians, red line shows the mean, and box limits indicate the 5^th^ and 95^th^ percentiles. Statistical significance was assessed via two-tailed t-test. **C**. Representative images of TMEM16F-HEK cells induced to express TMEM16F, treated with ionomycin, rAnxA5 and Ca^2+^ and stained with Cellmask Orange Actin (green) and rAnx A5 (red). Then the cells were treated with 10μM exogenous PC or PS. The images were taken10 min after adding exogenous lipids. **D**. Analysis of the experiments such as the one shown in **C** with an additional control of the cells not treated with any exogenous lipid. Average cortex detachment area per cell was measured in 4 independent experiments for more than 1,300 cells for each condition, normalized to detachment area/cell in control, and presented as box plots with center line showing the median, red line showing the mean, and box limits indicating the 5^th^ and 95^th^ percentiles. Statistical significance was assessed via two-tailed t-test.

As in fixed osteoclasts, fluorescent puncta of the PM-associated rAnx A5 on live osteoclasts, although much smaller than the Anx A5-enriched patches observed with α-Anx A5 Ab in fixed cells, correlated with reduced actin probe fluorescence, suggesting lower AC density beneath the PM at these sites (Fig. 3C). Thus, in both fixed and live osteoclasts, AC–PM detachment areas preferentially co-localized with PS-enriched regions at the cell surface.

To examine whether PS exposure is accompanied by weakening of AC-PM attachment in cells other than osteoclasts, we induced and activated scramblase expression and activity thereby promoting PS exposure in TMEM16F-HEK293 cells. As in differentiating osteoclasts, scrambling activation in TMEM16F-HEK293 cells led to the appearance of AC-PM detachment regions (Fig. 4A,B, Supplemental Movie 1). Application of rAnx A5 further promoted the detachment (Fig. 4B, Fig. S1). These data indicate that PS exposure and Anx A5 binding induce local AC-PM detachments, rather than vice versa: the detachments promoting local PS exposure and Anx A5 binding. Note that rAnx A5 application promoted the AC detachment only in cells with activated scramblase, as evidenced by the negligible detachment areas observed in control experiments where TMEM16F was either not induced or not activated (Fig. 4B).

The high efficiency of TMEM16F-mediated lipid scrambling ^53^ in the TMEM16F-HEK293 cells allowed us to efficiently deliver exogenous lipids to the inner PM leaflet by adding them to the outer leaflet. Exogenous NBD-tagged phosphatidylcholine (PC) and NBD-tagged PS associate with the cells in similar amounts (Fig. S2). However, only the influx of exogenous PC reduced the PS content in the inner leaflet. Our finding that adding PC to the medium promotes AC-PM detachment, compared to adding PS (Fig. 4C,D) supports the hypothesis that decreased PS concentration in the inner leaflet promotes AC detachment in scrambling cells.

In TMEM16F-HEK293 cells, to promote the AC-PM detachments, the scramblase had to be activated by raising intracellular Ca^2+^: no detachments were observed if the cells were not treated with Ca^2+^-ionophore (Fig. 4B). In differentiating osteoclasts, PS exposure and Anx A5 binding to the cell surface also depends on intracellular Ca^2+^. We loaded the differentiating osteoclasts with the cell-permeant Ca^2+^ probe Cal-520, AM and, as reported previously ^54^, observed Ca^2+^ oscillations (Supplemental Movies 2, 3). Earlier work has shown that these oscillations are critical for triggering the NFATc1-dependent transcriptional program at the onset of RANKL-induced osteoclast differentiation ^54,55^. Suppressing these oscillations and lowering intracellular Ca^2+^ levels using the cell-permeant chelator BAPTA inhibits these early stages of osteoclastogenesis. Importantly, while Ca^2+^ oscillations were initiated at relatively early stages of the differentiation following RANKL application, they persisted 3-4 days post-RANKL (^54^ and Supplemental Movies 2, 3, where Ca²⁺ flashes were observed in cells with externalized PS). If PS exposure at the fusion stage that follows NFATc1 signaling involves Ca^2+^ dependent scramblase, lowering intracellular Ca²⁺ should suppress synchronized fusion. Indeed, we found that BAPTA, AM robustly inhibits fusion observed after LPC removal (Fig. 5A,B). These data indicate that, in addition to its well-characterized role in the early stages of osteoclast differentiation ^54^, Ca^2+^ signaling promotes osteoclast fusion, possibly by activating and sustaining Ca^2+^ dependent scramblase activity. In further support of this hypothesis, osteoclasts with higher intracellular Ca^2+^ levels (as detected with Cal-520) showed rAnx A5 staining (Fig. S3), while lowering intracellular Ca^2+^ with BAPTA inhibited PS exposure, as detected by as rAnx A5 binding to the osteoclast surface (Fig. 5C,D).

**Fig. 5.**
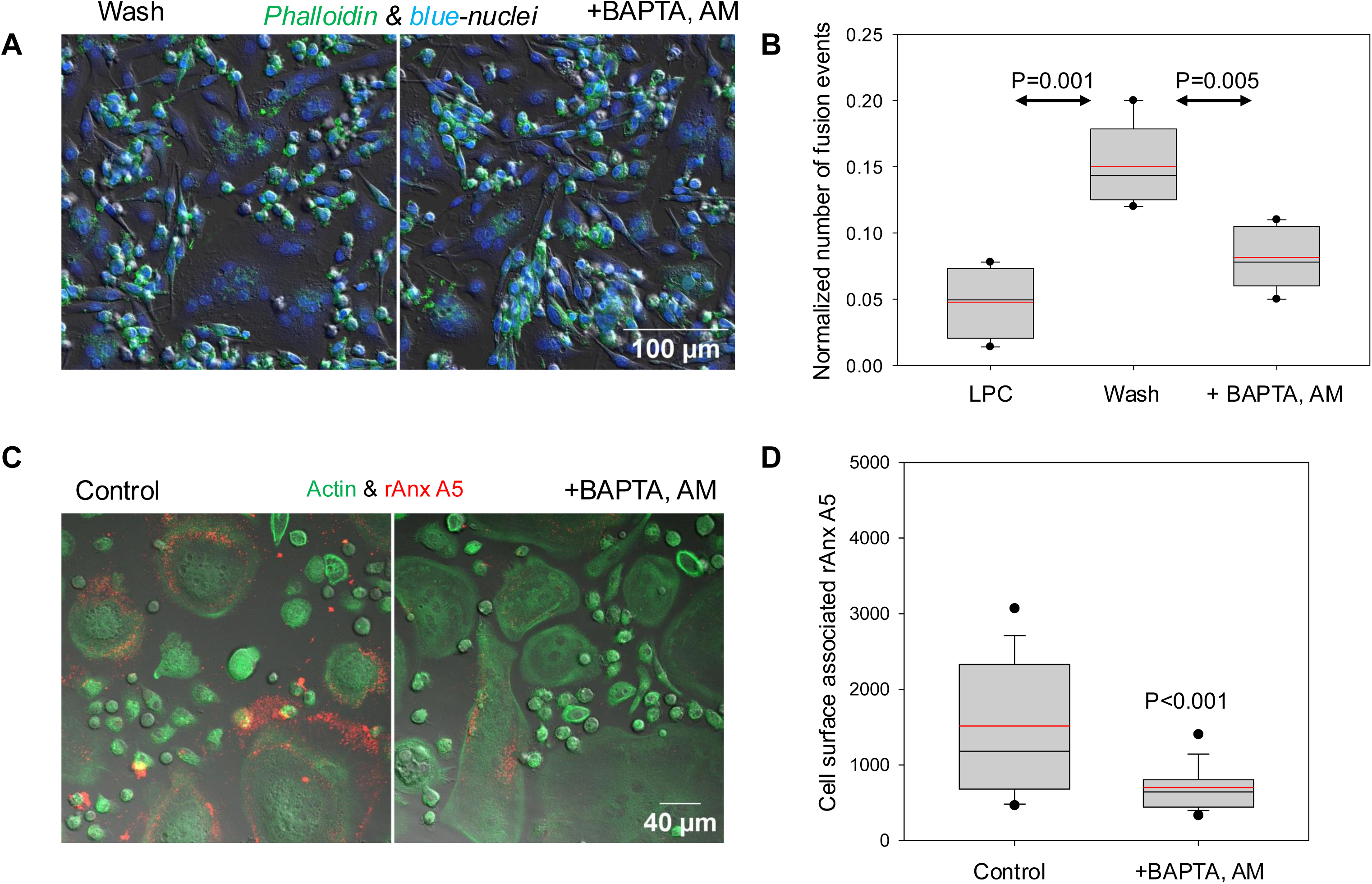
The dependence of fusion and PS exposure in osteoclasts on intracellular Ca^2+^ signaling. **A,B**. Cell-permeant BAPTA, AM inhibits osteoclast fusion. **A, B**. At 4 days post-RANKL application we placed the differentiating osteoclasts into LPC-containing medium for a 16-h time interval and then washed LPC out to allow fusion either in the presence of 25 μM BAPTA, AM or without it (“+ BAPTA, AM” and “Wash”, respectively). 2 hours after LPC removal we took images of the cells for off-line analysis. Representative fluorescence microscopy images (**A**) and quantification (**B**) of the numbers of cell fusion events with and without BAPTA, AM. “LPC” – fusion observed without removal of LPC for the cells not treated with BAPTA, AM. **C, D**. BAPTA, AM application to the differentiating osteoclasts at 4 days post RANKL application inhibits PS exposure. **C**. Representative fluorescence microscopy images of live differentiated osteoclasts incubated or not incubated with 20 μM BAPTA, AM (“+ BAPTA, AM” and “Control”, respectively) for 20 min at 37°C and then for 10 min at 37°C, stained with Cellmask Green Actin (green) and fluorescent rAnx A5 (red). **D**. Quantification of the experiments such as the one shown in **C** as intensity of cell surface Anx A5 labeling per field of view. **B, D.** Data from at least 10 random imaging fields for each condition in each of 3 independent experiments (>3, 000 cells, 60 fields for each condition, respectively), are presented as box plots with center line showing the median, red line showing the mean, and box limits indicating the 5^th^ and 95^th^ percentiles. Statistical significance was assessed via two-tailed t-test.

Our finding that fusion competence in differentiating osteoclasts is accompanied by PS exposure and Anx A5-dependent AC-PM detachments suggested that these detachments, and the resulting outward deformations of PM, play an important role in osteoclast fusion. To test this hypothesis, we applied NSC668394, a small molecule inhibitor of the phosphorylation of ERM proteins ^56^ and AC/PM attachment ^57^, to osteoclasts prior to LPC removal to allow synchronized fusion. We found that NSC668394 robustly promotes osteoclast fusion (Fig. 6A,B).

**Fig. 6:**
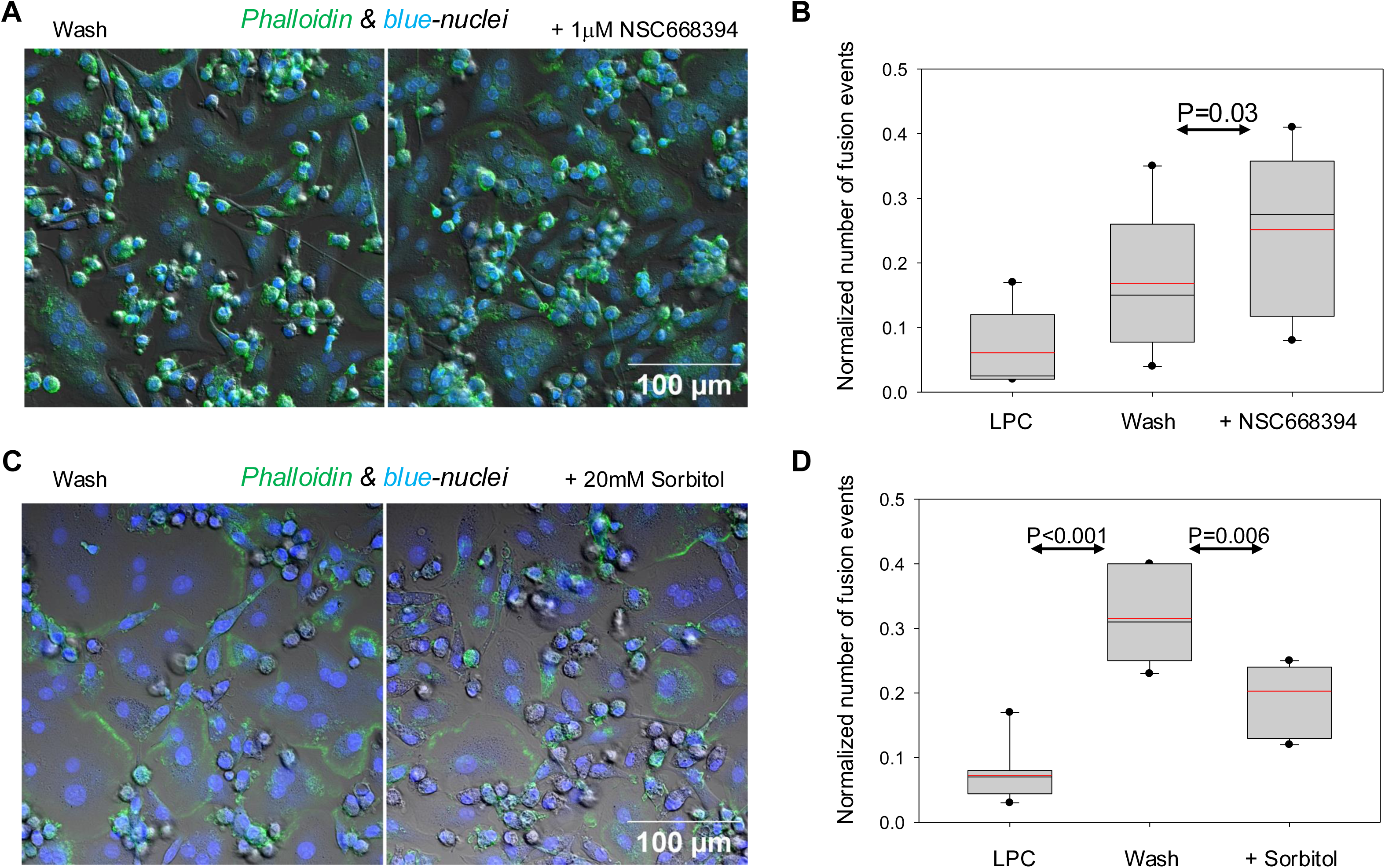
Osteoclast fusion depends on actin cortex-plasma membrane detachments and outward deformations of PM. **A, B**. Suppressing AC-PM attachment by ezrin phosphorylation inhibitor NSC668394 applied at the time of LPC removal promotes synchronized osteoclast fusion. A. Representative fluorescence microscopy images of differentiating osteoclasts 4 days post-RANKL application fixed 2 hours after LPC removal and stained with Phalloidin (actin filaments, green) and Hoechst (nuclei, blue) for the cells treated and not treated with 1 μM NSC668394 (“NSC668394” and “Wash”, respectively). **B**. Quantification of the numbers of cell fusion events in the presence of 0 (“Wash”) or 1 μM NSC668394. “LPC” – fusion observed without removal of LPC. **C,D**. Raising the tonicity of the medium by adding 20 mM sorbitol before LPC removal inhibited the efficiency of their synchronized fusion. **C**. Representative fluorescence microscopy images of osteoclasts 4 days post-RANKL application fixed 2 hours after LPC removal and stained with Phalloidin and Hoechst for the cells treated and not treated with 20mM sorbitol (“+ Sorbitol” and “Wash”, respectively. D. Quantification of the experiments such as the one shown in **C** with an additional control (“LPC”), where the cells not treated with sorbitol were kept in the presence of LPC. **B, D**. Data are pooled from 4 (NSC668394) and 3 (sorbitol) independent experiments (at least 10 random imaging fields all together more than 3,000 and 2,000 cells per condition in **B** and **D**, respectively). Data are presented as box plots with center lines showing the medians; red lines showing the mean, and box limits indicate the 5^th^ and 95^th^ percentiles. Statistical significance was assessed via two-tailed t-test.

We further hypothesized that AC-PM detachments promote fusion by facilitating local outward PM deformations driven by positive intracellular pressure ^26^. Thus, we expect that lowering intracellular pressure by placing osteoclasts in a hypertonic medium would inhibit fusion. Indeed, increasing medium tonicity by adding 20 mM sorbitol prior to LPC removal inhibited the efficiency of synchronized fusion (Fig. 6C,D). Note that, in addition to lowering intracellular pressure, hypertonic medium also reduces PM tension, and this is also expected to suppress fusion because tension drives opening and expansion of fusion pores ^58^.

The observation that osteoclast fusion is both accompanied and promoted by lipid scrambling and Anx A5– dependent AC–PM detachments motivated us to ask whether these detachments and detachment induced PM deformations are sufficient to induce cell-cell fusion. We co-incubated doxycycline-induced TMEM16F-HEK293 cells labeled with a cytosolic green cell tracker with cells labeled with cytosolic orange cell tracker, then applied ionomycin and rAnx A5 to promote PS exposure and AC detachment.

Thirty min later we examined the cultures for the cells co-labeled with both green and red probes. We considered the appearance of the cells co-labeled with both green and orange cell-trackers, as evidence of their fusion. We focused on contacting cells labeled with different trackers, where at least one cell was surface-labeled with Anx A5, indicating lipid scrambling, and examined whether these contacts yielded cells co-labeled with both trackers. Based on Wilson’s method ^59^, the probability of detectable fusion between TMEM16F-HEK293 was estimated to be below ∼0.6% (1 out of 994 contacts). These data indicated that, while lipid scrambling and cell-surface binding of Anx A5 promote osteoclast fusion, they are insufficient to induce fusion on their own, supporting earlier reports that fusogenic proteins are indispensable for cell-cell fusion ^60,61^.

In summary, Ca^2+^-dependent PS externalization—promoted by extracellular Anx A5 binding—lowers the PS content in the inner PM leaflet, inducing local AC-PM detachments. Promoting these detachments using an ezrin inhibitor enhances osteoclast fusion, whereas suppressing PM deformations and lowering membrane tension with sorbitol inhibits fusion. These findings support the hypothesis that AC restricts local PM deformations required for fusion.

### Computational model suggests a membrane bulging and an increase in membrane tension in the cortex detachment region

We propose that the AC-PM detachment events described above give rise to two interrelated phenomena within the membrane region disconnected from the cortical cytoskeleton: an increase in the lateral tension and a progressive bulging of the membrane surface. We suggest that the tension increase promotes the fusion pore formation and expansion, while the membrane bulging helps bridge the tens of nanometers wide gap established between the two membranes by the extracellular domains of the membrane proteins constituting cell-cell adhesion machinery.

To substantiate this proposal, we developed a theoretical model of the membrane conformation and dynamics in the region of the AC-PM separation. We used this model to computationally analyze the change in the membrane tension within the detachment region and the tendency of the detached membrane to bulge.

We describe the PM’s conformation and mechanical properties by the crumpling-stressing model, which was suggested in ^62,63^ and is based on the picket-fence concept of membrane organization ^64^. Briefly, the membrane is assumed to be subdivided into compartments with an average size of 200 nm ^64^ by a network of dense rows of transmembrane proteins, which are anchored to the underlying AC and serve as the compartment boundaries (Fig. 7A).

**Fig. 7:**
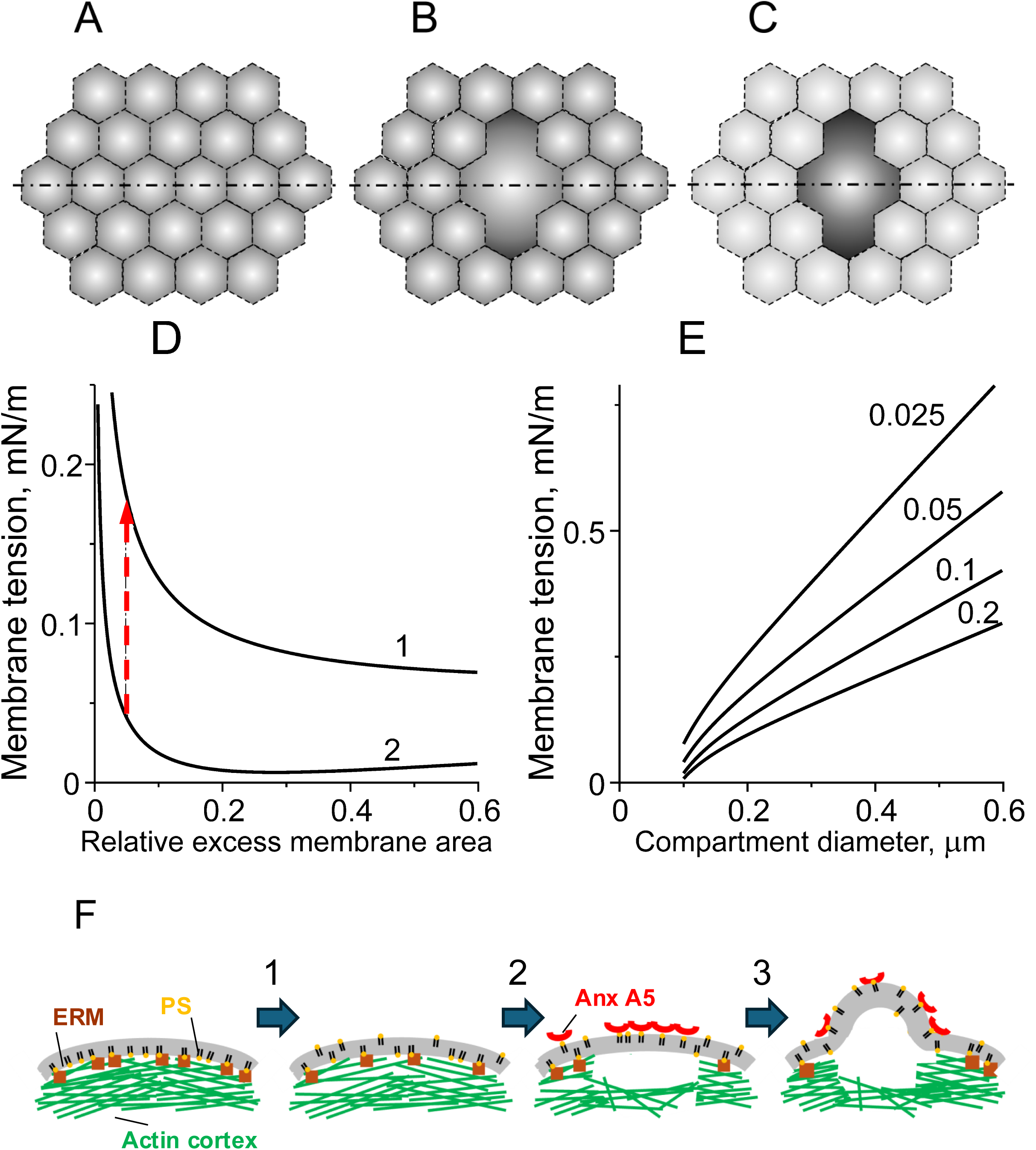
The proposed mechanism by which local detachment of actin cortex promotes fusion by facilitating pre-fusion membrane deformations with increased tension. **A, B, C**. Model of growth and bulging of cortex-organized membrane compartment. Top: top view of the PM; membrane compartments are represented by shaded grayscale regions. Bottom: side view of a system’s cross-section along the dash-dotted line. Compartments’ boundaries (dashed lines at the top) are formed by proteins (purple ellipses at the bottom) anchoring the PM (black) to AC (gray). **A**. The initial state of the system. **B**. The system immediately after the compartment merger resulting in the formation of a bigger compartment. **C**. The state of the system resulting from the redistribution of the membrane area into the newly formed big compartment accompanied by the swelling of the membrane bulge. **D**. The tension dependence on the relative excess membrane area for smaller (1) and bigger (2) compartments. The initial relative excess area was set to *β*_!_ = 0.05; the size of a small compartment (curve 1) was 0.2 μm; and that of the large compartment (curve 2) was 0.4 μm. Red arrow shows an increase in the PM tension caused by an increase in AC compartment size from 0.2 to 0.4 μm. **E**. The dependence of tension on the size of the compartment base for various compartment’s relative excess areas (indicated next to each curve). **D, E**. Membrane bending stiffness and the hydrostatic pressure difference were set to 10⁻¹⁹ J and 750 Pa, respectively. **F**. Hypothetical pathway from intracellular Ca^2+^ rise to PS-exposure to fusion-promoting deformation of plasma membrane, PM. In the initial state, the ERM (brown squares)-mediated connection between PS (yellow headed lipid)-enriched inner leaflet of PM & underlying actin cortex, AC (green lines) restricts membrane deformations. Transition 1: Lipid scramblase activated by intracellular Ca^2+^ rise releases anionic lipids, including PS, from the inner to outer leaflet of PM, and this results in a local detachment of a fraction of ERM and AC from the PM. Transition 2: Extracellular Anx A5 binds to cell surface PS and further decreases PS content in the inner PM leaflet and ERM- and AC-attachment. Transition 3: Bulging of the PM region disconnected from AC and local increase in tension in this PM region facilitates closer approach between fusing membranes and drives fusion pore formation.

The myosin-driven contraction of the AC - hence, of the associated compartment boundaries - results in the membrane bulging above the AC-PM interface, with the latter playing the role of an effective base plane of the membrane (Fig. 7A). The bulging occurs within each compartment and results in an overall crumpled membrane conformation. The intracellular pressure, *P*, pushes the membrane bulges, and sets the value of the membrane compartment’s tension. The exact value of the membrane tension changes along the surface of an inhomogeneously curved membrane of a bulge ^63,65^. For convenience, in the following, we characterize the PM bulge’s tension by its constant component, *γ*, whose physical meaning is the tension in a virtual flat membrane, which would be in mechanical equilibrium with the bulge’s membrane if connected to it. Considering the membrane monolayers to be non-stretchable, the value *γ* can be related to the chemical potential of the lipid material, *μ*, by *γ* = *μ*_0_/*a* − *μ*/*a*, where *a* is a constant in-plane area per lipid and *μ*_!_ is the lipid chemical potential of a flat membrane upon a vanishing tension. For brevity, we refer to this constant component, *γ*, as, simply, the tension.

We assume that in the initial state preceding the AC-PM detachment, all the membrane compartments are similar and characterized by the same values of the initial tension *γ*_“#_, the membrane area stored within a compartment, *A*, and the area of the compartment’s projection on the base plane, *A*_$_, referred to below as the base area. To quantify the extent of the membrane bulging within a compartment, we use the compartment’s relative excess area, 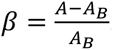.

We model the local AC-PM detachment as the vanishing of proteinaceous boundaries between several adjacent compartments, resulting in the effective merger of these compartments into a larger one (Fig. 7A-C). The larger the detachment area, the larger the amount of the dissociated boundaries and, hence, the larger the stored membrane area, *A*, and the base area, *A*_$_, of the newly formed compartment. Immediately after the compartments’ merger, the relative excess area, *β*, of the new compartment is equal to that of the initial compartments.

Our goal is to make predictions on the evolution of the membrane tension in the newly formed big membrane compartment and the membrane area redistribution between this compartment and the surrounding compartments that remained unaffected by the AC-PM detachment (Fig. 7A-C). For this purpose, we computationally analyzed the relationships between the membrane tension, *γ*, the stored membrane area, *A*, and the base projection area, *A*_$_, of a membrane compartment upon a fixed value of the intracellular pressure, *P*. The equations and techniques used in these computations are described in detail in ^62,63,66^ and briefly outlined in Supplemental Appendix A.

The essence of the obtained computational results presented in (Fig. 7D,E) is that the compartment’s tension, *γ*, is predicted to rise upon an increase of the compartment’s dimension and, hence, upon a merger of a few compartments into a larger one as a result of the AC-PM detachment. Fig. 7D presents the tension dependence on the compartment’s excess area, *β*, for two different values of the base projection size, +*A*_$_, and demonstrates that the tension is larger in a larger compartment for all values of *β*. Fig. 7E illustrates a monotonic increase in tension with increase in the base projection size, +*A*_$_, for different values of the compartment’s relative excess area, *β*, within a range suggested in ^63,67^ with membrane bending stiffness and the hydrostatic pressure difference set to 10⁻¹⁹ J ^68^ and 750 Pa ^26^, respectively.

An important consequence of the rise of the compartment’s tension with the compartment’s size is a prediction that the merger of several compartments must initiate the membrane area redistribution into the emerged bigger compartment at the expense of the area stored in the surrounding compartments, which must lead to swelling of the big compartment’s membrane bulge. Indeed, due to the membrane lateral fluidity, the membrane material must flow along the membrane plane in the direction opposite to the in-plane gradient of the chemical potential, *μ*, and, hence, along the in-plane gradient of the tension, *γ*. Therefore, the merger-related instantaneous increase in the compartment’s tension, *γ*, compared with the initial tension, *γ*_“#_, existing in the neighboring unperturbed compartments, must drive the membrane area redistribution from the surrounding into the newly created bigger compartment and the related swelling of its membrane bulge. These processes are mediated by the inter-compartment in-plane flow of the membrane material, which is driven by the tension difference, Δ*γ* = *γ* – *γ_in_*, and limited by the permeability of the compartment boundaries ^62,63^.

It must be emphasized that the above analysis and reasoning are valid only for the time point right after the compartments’ merger. Starting at this initial time point, the process of the area redistribution is predicted to gradually reduce the tension difference between the newly formed and the surrounding compartments, Δ*γ*. The redistribution process and the related bulge swelling would stop when the tension reaches a full equalization between all the compartments, including the bigger one formed through the compartment merger. This equalization is predicted to result in a new tension level, *γ_fin_*, which is larger than the initial one, *γ_fin_* > *γ_in_*, but smaller than the instantaneous tension, *γ*, generated in the bigger compartment right after the merger event.

In summary, the bulging of the PM membrane region at the AC-PM detachment site is subjected to an elevated lateral tension suggesting that AC detachment facilitates both the local membrane swelling mediating the mutual approach of the opposing membrane bilayers and the fusion pore formation.

## DISCUSSION

While different cell fusion processes utilize different specialized proteins (reviewed in ^1^), diverse fusions, including the formation of multinucleated osteoclasts, are accompanied by intracellular calcium signaling, non-apoptotic exposure of PS at the surface of fusion-committed cells and the cell surface binding of extracellular Anx A5 (reviewed in ^12^). Our findings suggest that each of these hallmarks of cell fusion represents a step in a conserved pathway that leads to a localized weakening of the AC-PM attachment, facilitating the formation of tight pre-fusion PM bilayer contacts and fusion intermediates (Fig. 7F). Below, we discuss the distinct stages of this pathway.

At the early stages of osteoclast differentiation triggered by RANK/RANKL interactions, cytoplasmic Ca^2+^ oscillations upregulate NFATc1, a master regulator of osteoclastogenesis ^6^. Here, we report that downstream of Ca^2+^-NFATc1 signaling, cell fusion, a relatively late stage of osteoclast formation, also depends on intracellular Ca^2+^. Fusion inhibition by lowering intracellular Ca^2+^ in ready-to-fuse cells was accompanied by reduced PS exposure reflected in a lowered rAnx A5-cell surface binding. Inhibition of fusion by BAPTA, AM argues an alternative interpretation that the decrease in rAnx A5 binding reflects an increase in the amount of endogenous Anx A5 at the cell surface. The molecular machinery responsible for disrupting the asymmetric distribution of phospholipids between the inner and outer PM leaflets and delivering PS to the surface of differentiating osteoclasts remains to be clarified. Some earlier studies suggested that PS exposure in osteoclasts involves the floppases ABCC5 and ABCG1 ^69^ and/or the caspase 8-activated scramblase XKR8 ^11^. However, the intracellular Ca^2+^ dependence of the PS externalization shown here, along with the inhibition of both PS exposure and osteoclast fusion by CaCCinh-A01, an inhibitor of calcium-activated chloride channels and TMEM16 family scramblases, reported in an earlier study ^8^, suggests the involvement of a TMEM16 scramblase.

While binding of rAnx A5 to cell-surface PS is widely used to detect apoptotic cells and cells dying via other pathways ^70^, binding of endogenous extracellular Anx A5 to PS at the surface of living cells plays an important role in numerous physiological and pathological processes ^71^. Extracellular Anx A5 has been proposed to influence cell physiology by interacting with different partner proteins, including S100A4 ^8^ and La protein ^14^; competing with some of the PS binding proteins such as prothrombin ^72^; self-assembling into two-dimensional arrays on membrane surfaces ^73^; generating negative (concave) curvature of the membrane ^74^; and inducing phase transitions in the underlying lipid ^75^. Here, we found that Anx A5 binding to PS in the outer leaflet of PM with activated scramblase may affect cell physiology by altering the trans-leaflet distribution of PS. Lipid scrambling may, at least locally, halve the concentration of PS in the inner leaflet of PM by allowing PS molecules to redistribute to the outer leaflet initially depleted of this lipid species. High-affinity binding of the extracellular Anx A5 to cell surface PS, with a dissociation constant in the nanomolar range (K_d_ < 0.1 nM ^76^ and ∼15 nM ^77^), is expected to sequester additional PS molecules from the inner to the outer PM leaflet, effectively acting as a “PS trap”.

PS in the inner leaflet of the PM plays a critical role in recruiting and regulating many cytosolic proteins, including those involved in PM dynamics and rearrangements ^78,79^ , as well as ERM family proteins that link the actin cortex to the PM. The transition of ERM proteins from a cytoplasmic, closed conformation to an activated, open conformation, serving as linkers between the PM and the contractile actomyosin network, depends on PS and PI(4,5)P₂, another anionic lipid primarily localized to the inner PM leaflet ^31–34^. The surface concentration of PI[4,5]P_2_ regulates ERM recruitment, and PS synergistically promotes the formation of large-scale membrane-associated actin networks with myosin II motors ^34^. We found that Ca^2+^ dependent lipid scrambling in differentiating osteoclasts reduces both PS and PI[4,5]P_2_ content in the inner leaflet of PM. This scrambling-induced loss of PI[4,5]P_2_ is consistent with an earlier report documenting the scramblase dependent appearance of this lipid in the outer PM leaflet ^38^. Moreover, because Anx A5 binds to PI[4,5]P_2_ ^80^, Anx A5 binding to cell-surface PI[4,5]P_2_ is expected to further reduce the concentration of this lipid in the inner leaflet. Our finding that activation of lipid scrambling leads to a loss of intracellular PM-associated PI(4,5)P2 is consistent with two earlier studies, in which scrambling and blebbing were induced by treating cells with dithiothreitol and paraformaldehyde ^81,82^. Note that in these studies, the depletion of PI(4,5)P2 from the inner PM leaflet was attributed to its degradation rather than redistribution to the outer leaflet.

Fusion depends on membrane deformations. While, as recently suggested, lipid scrambling may directly facilitate bulging of PM by lowering the bending rigidity of the lipid bilayer ^82^, our findings suggest that in differentiating osteoclasts and in TMEM16F-HEK293 cells, scrambling weakens AC-PM attachment that plays a critical role in controlling PM deformations. Our findings that extracellular Anx A5 promotes both AC-PM detachment and fusion, and that ERM inhibitor suppresses both AC-PM attachment and fusion suggests that AC detachments are involved in osteoclast fusion. This conclusion is consistent with two recent studies.The first reports that decreasing AC-PM attachment by depleting or inhibiting ERM protein moesin promotes formation of tunneling nanotubes and multinucleated osteoclasts ^20^. The second study documents downregulation of the ERM protein ezrin in differentiating Raw 264.7 cells and finds that reduced ezrin-mediated AC-PM attachment is a prerequisite for osteoclast fusion ^44^.

The dependence of cell fusion on AC–PM detachments may also explain an earlier observation that the small GTP-binding protein Rnd3/RhoE, recruited to AC-free patches of the PM to suppress cortex reassembly ^83^, promotes myoblast fusion ^84^. It is important to note that the fusion-restricting contributions of the AC during membrane fusion, discussed here, are distinct from the many roles of the actin cortex in earlier stages of multinucleated cell formation, including cell–cell adhesion ^85^ and the post-fusion syncytial cell reshaping ^86^, and the fusion-promoting contributions of the tightly packed, parallel arrays of actin filaments in actin bundles ^4^.

Local detachment of the PM from the underlying AC initiates outward bulging of PM driven by the contraction of the actomyosin cytoskeleton ^27^. This bulging gives rise to blebs ^87^, initiates actin bundle-propelled cell protrusions ^27^ and facilitates the shedding of protrusion-derived extracellular vesicles ^88^. AC detachment is expected to promote local bulging of PMs that bring the membrane bilayers of the two PMs closer to each other than the 25 nm distances characteristic for cell-cell contacts, supported by the E-cadherin adhesion machinery ^89^, involved in the formation of multinucleated osteoclasts ^90^. An elevated fraction of disordered domains in the PM regions detached from the AC ^91^ may further facilitate close approach and fusion by easing protein displacement from the pre-fusion tight contacts between protein-depleted patches.

Our theoretical model suggests that a local AC-PM detachment promotes fusion by generating outgoing deformation of PM and increasing membrane tension. We consider PM to be subdivided into compartments by a network of rows of transmembrane proteins anchored to AC. Within each compartment, the membrane is assumed to bulge because of AC contraction and be stressed by the intracellular pressure, which pushes the membrane and generates the membrane tension. Our computational analysis shows that an effective merger of several compartments into a bigger one, because of the AC-PM detachment, results in a local increase in the membrane tension. This increase can be comparable with the initial tension. While elevated PM tension can inhibit formation of the early hemifusion intermediates ^92^, tension increase is expected to promote most energy demanding stages of fusion pathway: opening and expansion of a nascent fusion pore^58^. Note that the predicted growth of the membrane tension is in apparent contradiction with the recent paper ^44^ suggesting that the AC dissociation from the PM decreases rather than increases the membrane tension. The conclusion of ^44^ is based on the measured decrease in the tether pulling force, which includes two independent contributions, those of the membrane tension and the energy of the AC-PM attachment ^93^. It is our understanding that in the system explored in ^44^ the decrease in the attachment energy overcomes the increase in the membrane tension, which results in an observed reduction of the tether-pulling force. Note that lowering PM tension by placing the cells in hypertonic medium in our experiments inhibited osteoclast fusion.

In conclusion, our findings uncover a bidirectional signaling pathway in which inside-out steps —including an intracellular Ca^2+^ dependent activation of a lipid scramblase that delivers PS to the surface of the cell — are followed by outside-in steps, including Anx A5-dependent depletion of PS from the inner PM leaflet, which leads to a local detachment of the AC. We uncovered this signaling pathway in fusion-committed osteoclasts, and we propose that AC detachment promotes the cell fusion stage of osteoclastogenesis, as well as other fusion processes, by facilitating the necessary PM deformations. However, weakening the AC-PM attachment via lipid scrambling and Anx A5-PS interactions at the surface of the cells may also play an important role in other physiological contexts. The extracellular environment may contain substantial amounts of Anx A5 ^71^; and transient PS exposure in living cells, in addition to being a shared hallmark of diverse cell-cell fusion processes (reviewed in ^12^), is observed in the immune response ^94^; neuronal regeneration and degeneration ^95^, and in cancer cells in metastases ^96,97^. Furthermore, beyond their role in the AC-PM attachment, both PS and PI(4,5)P2 in the inner PM leaflet regulate cell physiology by controlling PM recruitment of numerous cytosolic proteins with cationic domains or PS-binding motifs ^98^ or PI(4,5)P2-binding motifs such as pleckstrin homology (PH) domains ^99,100^. Redistribution of these proteins from the PM to the cytosol, triggered by lipid scrambling and Anx A5–dependent depletion of anionic lipids, is expected to profoundly alter many associated cellular properties. A better understanding of the cascade of events within this emerging inside-out/outside-in signaling pathway will likely uncover novel mechanisms that coordinate diverse aspects of cellular responses in various biological contexts.

## Author contributions

E.L., J.M.W. and L.V.C. designed experiments; M.M.K. and A.K.T conceived and performed the theoretical studies. E.L., J.C., M.R. and G.K. performed research; E.L., K.M. and L.V.C. analyzed data; and L.V.C., A.K.T. and M.M.K. wrote the paper.

## Acknowledgements

We wish to thank the Criss Hartzell Laboratory at the Emory University for sharing the HEK-TMEM16F cell line and the National Institutes of Health Department of Transfusion Medicine for isolating the monocytes used in this study. The research in L.V.C. laboratory was supported by the Intramural Research Program of the Eunice Kennedy Shriver National Institute of Child Health and Human Development, National Institutes of Health. MMK acknowledges support by the Israel Science Foundation (Grant No. 1994/22) and holds Joseph Klafter Chair in Biophysics. We used ChatGPT (https://chatgpt.com/) to assist with grammar and stylistic refinement of the final manuscript draft. Prompts included “help check for typos and grammatical errors.” All outputs were carefully reviewed and edited by the authors to ensure accuracy.

## METHODS

### Reagents

We purchased human M-CSF and RANKL from Cell Sciences (catalog #CRM146B and #CRR100B, respectively). LPC (1-lauroyl-2-hydroxy-sn-glycero-3-phosphocholine, #855475); NBD-PC (1-palmitoyl-2-{6-[(7-nitro-2-1,3-benzoxadiazol-4-yl)amino]hexanoyl}-sn-glycero-3-phosphoserine (ammonium salt) in chloroform, catalog # 810192C) and NBD-PS (1-palmitoyl-2-{6-[(7-nitro-2-1,3-benzoxadiazol-4-yl)amino]hexanoyl}-sn-glycero-3-phosphocholine, catalog # 810130C) from Avanti Polar Lipids. Hoechst 33342 and Alexa Fluor Plus Alexa555 phalloidin were purchased from Invitrogen (catalog #H3570 and A30106, respectively). We labeled F-actin in live cells with CellMask Green Actin or Cellmask Orange Actin tracking stains (Invitrogen, catalog # A57246 and #A57244, respectively) or with SiR-actin Kit, (Cytoskeleton, catalog # Cy-SC001). Membrane stain FM 4-64 Dye (N-(3-Triethylammoniumpropyl)- 4-(6-(4-(Diethylamino) Phenyl) Hexatrienyl) Pyridinium Dibromide) was purchased from Thermo Fisher Scientific (Catalog # T13320). We purchased membrane-permeant Green CMFDA cell tracker and orange CMRA Cell Tracker from ThermoFisher Scientific (#C7025 and #C34551, respectively).

We used constructs for expressing intracellular probes for PS (Lact-C2-GFP, Plasmid #22852 ^101^, a gift from Sergio Grinstein), and PI(4,5)P2 (PIP2-CFP(UST) sensor based on the PH domain of PLCd1, Plasmid #170357, a gift from Kees Jalink. Ezrin Inhibitor, NSC668394 was purchased from Sigma Aldrich, catalog #341216. Ionomycin was purchased from Tocris Bioscience, catalog *#*1704. Cell-permeant chelator BAPTA, AM and calcium-sensitive fluorescent dye Cal-520 AM that we used to evaluate changes in intracellular calcium were purchased from Invitrogen (catalog # B6769) and AAT Bioquest (catalog #21135), respectively. We purchased unlabeled recombinant A5 (rAnx A5) from BD Pharmingen Cat 51-65871A and recombinant fluorescent Annexin V Conjugates CF568 from Biotium, catalog #29010. D-Sorbitol was purchased from Sigma-Aldrich (catalog # W302902). Doxycycline hyclate was purchased from Sigma (catalog #D9891-5G). Secondary Anti-mouse IgG (H+L), F(ab’)2 Fragment (Alexa Fluor® 647 Conjugate) was purchased from Cell Signaling (catalog #4410S).

### Osteoclasts

Elutriated monocytes from healthy donors were obtained through the Department of Transfusion Medicine at National Institutes of Health under protocol 99-CC-0168 approved by the National Institutes of Health Institutional Review Board. Research blood donors provided written informed consent and blood samples were de-identified prior to distribution, Clinical Trials Number: NCT00001846. We also used elutriated monocytes from healthy donors obtained through Elutriation Core Facility, University of Nebraska Medical Center, informed consent was obtained under an Institutional Review Board approved protocol for human subject research 0417-22-FB. Primary human osteoclasts were derived as described in ^14^. Briefly, elutriated monocytes were added to complete media [α-MEM (Gibco) +10% FBS (Gibco)+ 1x penicillin/streptomycin/glutamate (Gibco)] supplemented with 100 ng/ml recombinant M-CSF and plated at 1×10^6^ cells/ml for 6 days. Next, the cells were placed into complete medium supplemented with 100 ng/ml recombinant M-CSF and 100 ng/ml recombinant RANKL and in 3 to 5 days generated multinucleated osteoclasts.

Stated concentrations of unlabeled rAnx A5 were applied to differentiating osteoclasts from 0.5 mg/ml stock solution. To stain osteoclast surface PS and polymerized/ filamentous actin (F-Actin) in live cells we placed differentiating osteoclasts in 35mm Ibidi dishes into 140 mM NaCl, 5 mM KCl, 5 mM MgCl_2_, 5 mM CaCl_2_, 10mM Tris buffer for 5 min. Then we washed the cells twice with warm complete medium, labeled them with CellMask Green Actin diluted 1:1000 and placed them into 1 ml of complete medium supplemented with 5 μl of 50 μg/mL stock solution of fluorescent rAnx A5. In time-lapse experiments on osteoclasts, we added 5 μl of fluorescent rAnx A5 from 50 μg/ml stock solution to the warm complete medium already under microscope.

To lower intracellular Ca^2+^ (Fig. 5C,D), we treated differentiating osteoclasts with 20 μM BAPTA, AM for 20 min at 37°C in complete medium, and, when needed, applied fluorescent rAnx A5 for 10 min at 37°C immediately before microscopy.

In some experiments, to evaluate Ca^2+^ signaling, we loaded osteoclasts with intracellular Ca^2+^ probe by incubating the cells in α-MEM supplemented with 5μM Cal-520, AM for 20 min at 37°C and then for 10 more minutes in the presence of rAnx A5.

### TMEM16F-HEK293 cells

We propagated TMEM16F-HEK293 cells ^51^ in DMEM (Gibco) containing 10% heat-inactivated fetal bovine serum and supplemented with penicillin-streptomycin (1%) at 37°C and 5% CO_2_. To induce exogenous TMEM16F expression, we plated the cells on 35mm Ibidi plates and incubated them with 1μM doxycycline for 16 h. Then, to trigger lipid scrambling we placed the cells for 5 min in 140 mM NaCl, 5 mM KCl and 5 mM MgCl_2_, 10mM Tris buffer (buffer A) supplemented with 1 μM ionomycin and either cell-permeable CellMask Actin Tracking Stain (Green or Orange) or SiR-actin Kit, both used for live imaging according to the manufacturer’s instructions (diluted 1:1000 and at a 1 μM final concentration, respectively). We washed the cells twice with the warm buffer A. Then we added to 1mL of warm buffer A bathing the cells 5 μl of 50 μg/ml stock solution of fluorescent rAnx A5 and CaCl_2_ to a final concentration of 5mM. Images of live cells were taken for analysis after 5 min incubation at the room temperature. In time lapse experiments, we added Ca^2+^ to a final 5mM concentration already under microscope. In Fig. 3B, to remove noise, images were smoothed with gaussian filter (1px radius). To emphasize structures of interest images were linearly contrasted using “Brightness/Contrast” tool to remove cytosolic background, out of focus signal and to reveal weakly fluorescent Annexin spots. Quantitative analysis of CellMask Actin fluorescence in Fig. 3C was performed using unmodified images.

To treat TMEM16F-HEK293 cells with exogenous NBD-PS and NBD-PC, 16 h after application of 1μM doxycycline, we placed the cells into buffer A supplemented with 1 μM ionomycin and CellMask Orange Actin and exogenous lipids from their 1mg/ml stock solutions in ethanol to a final 10μM concentration. After 5 min incubation at 37° C, we added 5 μl of fluorescent rAnx A5 and, at the same time, triggered lipid scrambling by adding CaCl_2_ to a final concentration of 5mM. Photos for analysis were taken after 5 min incubation of the cells at 37°C.

In some experiments, we transfected TMEM16F-HEK293 cells to express intracellular probes for PS and PI(4,5)P2. To express these probes: Lact-C2-GFP probe for PS and PLCδ1-based PIP2-CFP(UST) probe for PI(4,5)P2 in TMEM16F-HEK293 cells, we transfected the cells using lipofectamine 2000, 5μg DNA per plate for 2h, and then applied doxycycline for 16h to simultaneously induce TMEM16F and lipid probe expression. Scrambling was activated as described above. To evaluate changes in PS and PI(4,5)P2 content in the inner monolayer of PM we divided average per pixel fluorescence under randomly selected linear areas along the PM of the cell (>4 regions/per cell and >45 cells, for each condition) by average per pixel fluorescence of the similar areas randomly selected in cell cytoplasm.

### Fluorescence microscopy

To analyze osteoclast fusion efficiency, we washed cells with PBS and then rapidly fixed with warm, freshly prepared 4% formaldehyde in PBS at 37° °C. The cells were subsequently washed with PBS. We incubated them for 5 min in 0.1% Triton X100 in PBS. We stained osteoclast cytoskeletal boundaries with Phallodin-Alexa Fluor, 1:2000 dilution) and nuclei with Hoechst 33342 (1:5000 dilution). for 1 hr in complete staining buffer (PBS + 5% FBS + 0.1% Triton X100) before a final PBS wash prior to imaging.

For immunofluorescence analysis of the fixed cells, we incubated them in PBS supplemented with 10% FBS for 10 min at room temperature to suppress non-specific binding. Then, cells were incubated with primary antibodies to Anx A5 for 1 hr in PBS supplemented with 10% FBS. After 5 washes in PBS, we incubated the cells with fluorescent secondary IgG (1:1000 dilution) for 1 hr in PBS supplemented with 10% FBS (Anti-mouse IgG (H+L), F(ab’)2 Fragment (Alexa Fluor® 647 Conjugate) #4410, 1:1000 dilution) and then washed 5 times with PBS prior to imaging.

Images were captured on a Zeiss LSM 800, confocal microscope using a C-Apochromat 63x/1.2 water immersion objective lens and analyzed using Fiji/ImageJ’s open-source image processing package v.2.1.0/1.53c.

### Synchronization of osteoclast fusion

We synchronized cell fusion as in ^14^. Briefly, 3-4 days post RANKL application, osteoclast medium (α-MEM with 10% Fetal Bovine Serum (FBS), penicillin-streptomycin-L-glutamine and 100 ng/ml M-CSF, 100 ng/ml RANKL) was refreshed with media supplemented with 100 ng/ml M-CSF, 100 ng/ml RANKL and 350 μM lauroyl LPC. 16 h later, LPC was removed by 2 washes with fresh culture medium without LPC, RANKL and M-CSF supplemented or not supplemented with reagent under consideration (rAnx A5 in the stated concentrations or 20μM BAPTA, AM or 1μM NSC668394 or 20 mM sorbitol). The cells were allowed to fuse for 2 hours at 37°C.

Osteoclast fusion was scored as described earlier ^14^. In brief, we fixed the cells and labeled cell nuclei with Hoechst 33342 and actin filaments with phalloidin. Images of the 10 randomly selected fields of view were captured for off-line analysis. We evaluated the efficiency of osteoclast fusion by counting the numbers of syncytia (cells with two or more nuclei) and the numbers of nuclei in each syncytium.

### Cell fusion assay for TMEM16F-HEK293 cells

We labeled the cells in one dish with CellTracker Green and in another with Orange CMRA CellTracker by adding 2 μl of the CellTracker Green (or Orange CMRA CellTracker) stock solution (50 μg of the probe in 20 μl of DMSO) to 2 ml of PBS and incubated the cells in this medium for 15 min at 37°C. After two washes with the medium, we placed the cells into complete medium for 30 min at 37°C. Then we lifted the cells with 1 ml of trypsin-EDTA 0.05% (1 min at room temperature) and plated at Ibidi 35 mm dishes. After 2h incubation, we replaced the medium and induced TMEM16F expression by applying doxycycline overnight. TMEM16F scramblase was activated as described above. After 5 min at 37°C (with Ca^2+^ and fluorescent rAnx A5), we replaced the medium with fresh complete medium for 30 min and scored for the appearance of double labeled cells. For an upper estimate of possible fusion efficiency, we determined the binomial proportion confidence interval using the Wilson score method, and report the estimate as the upper limit of the 95% confidence interval ^59^.

### Cell fusion quantification

Osteoclast fusion efficiency was evaluated as the number of fusion events between cells as described in ^102^. The number of cell-to-cell fusion events required to generate syncytium with N nuclei is always equal to N-1. We calculated the fusion number index as Σ (Ni − 1) = N_total_ − N_syn_, where N_i_ = the number of nuclei in individual syncytia and N_syn_ = the total number of syncytia. We normalized the number of fusion events to the total number of nuclei (including unfused cells) to control for small variations in cell density from dish to dish and image to image.

### Quantification of the changes in PM-associated actin cortex density

We evaluated the local changes in the density of PM associated actin cortex in live differentiating osteoclasts in the presence of rAnx A5 4 days post RANKL application. We focused on regions with clearly identifiable AC seen as continuous ridge-like structures labeled with CellMask Green Actin. Within those regions, we compared average per pixel intensity of CellMask Green Actin for all areas that were stained with fluorescent rAnx A5 with the sample of Anx A5 negative areas located in the proximity to the rAnx A5 positive spots.

In TMEM16F-HEK293 cells, we quantified actin cortex detachment areas for cells in randomly selected fields of view by measuring for each of the cells the total area of the space between PM labelled with FM 4-64 and actin cortex labelled with CellMask Green Actin probe or, in the experiments with NBD-tagged exogenous lipids, with CellMask Orange Actin probe.

### Quantification and statistical analysis

Each set of experiments for each graph presented here was repeated on at least three occasions. Due to the inherent variability in the derivation of primary human monocytes from different donors, the precise time course of the differentiation and the efficiencies of fusion varied between experiments. However, the reported differences between groups were observed in each experiment. The data from multiple fields of view examined over at least three independent experiments were analyzed using a parametric two-tailed *t*-test to determine significance. Data are presented in box-and-whisker plots where the center lines show the median, the red lines the mean, the box the 25^th^-75th percentiles, whiskers above and below the box indicate the 90th and 10th percentiles, solid circles show 5th/95th percentile. These statistical analyses were performed using the SigmaPlot15 software. Normally distributed data were analyzed using the unpaired t test. When the data were not normally distributed or failed the equal variance test, we used the Mann-Whitney rank sum test instead. The P values for each statistical comparison are defined in each figure.

## Notes

### Competing Interest Statement

The authors have declared no competing interest.

