## Supplemental information for "PHOSPHATIDYLSERINE EXPOSURE AND EXTRACELLULAR ANNEXIN A5 WEAKEN THE ACTIN CORTEX IN OSTEOCLAST FUSION"

The essence of the obtained computational results presented in (Fig. 7D,E) is that the compartment's tension, $\gamma$, is predicted to rise upon an increase of the compartment's dimension and, hence, upon a merger of a few compartments into a larger one as a result of the AC-PM detachment. Fig. 7D presents the tension dependence on the compartment's excess area, $\beta$, for two different values of the base projection size, $\sqrt{A_B}$, and demonstrates that the tension is larger in a larger compartment for all values of $\beta$. Fig. 7E illustrates a monotonic increase in tension with increase in the base projection size, $\sqrt{A_B}$, for different values of the compartment's relative excess area, $\beta$, within a range suggested in [63,67] with membrane bending stiffness and the hydrostatic pressure difference set to $10^{-19}$ J [68] and 750 Pa [26], respectively.

An important consequence of the rise of the compartment's tension with the compartment's size is a prediction that the merger of several compartments must initiate the membrane area redistribution into the emerged bigger compartment at the expense of the area stored in the surrounding compartments, which must lead to swelling of the big compartment's membrane bulge. Indeed, due to the membrane lateral fluidity, the membrane material must flow along the membrane plane in the direction opposite to the in-plane gradient of the chemical potential, $\mu$, and, hence, along the in-plane gradient of the tension, $\gamma$. Therefore, the merger-related instantaneous increase in the compartment's tension, $\gamma$, compared with the initial tension, $\gamma_{in}$, existing in the neighboring unperturbed compartments, must drive the membrane area redistribution from the surrounding into the newly created bigger compartment and the related swelling of its membrane bulge. These processes are mediated by the inter-compartment in-plane flow of the membrane material, which is driven by the tension difference, $\Delta\gamma = \gamma - \gamma_{in}$, and limited by the permeability of the compartment boundaries [62,63].

It must be emphasized that the above analysis and reasoning are valid only for the time point right after the compartments' merger. Starting at this initial time point, the process of the area redistribution is predicted to gradually reduce the tension difference between the newly formed and the surrounding compartments, $\Delta\gamma$. The redistribution process and the related bulge swelling would stop when the tension reaches a full equalization between all the compartments, including the bigger one formed through the compartment merger. This equalization is predicted to result in a new tension level, $\gamma_{fin}$, which is larger than the initial one, $\gamma_{fin} > \gamma_{in}$, but smaller than the instantaneous tension, $\gamma$, generated in the bigger compartment right after the merger event.

Fig. 7: **The proposed mechanism by which local detachment of actin cortex promotes fusion by facilitating pre-fusion membrane deformations with increased tension. A, B, C**. Model of growth and bulging of cortex-organized membrane compartment. Top: top view of the PM; membrane compartments are represented by shaded grayscale regions. Bottom: side view of a system's cross-section along the dash-dotted line. Compartments' boundaries (dashed lines at the top) are formed by proteins (purple ellipses at the bottom) anchoring the PM (black) to AC (gray). **A**. The initial state of the system. **B**. The system immediately after the compartment merger resulting in the formation of a bigger compartment. **C**. The state of the system resulting from the redistribution of the membrane area into the newly formed big compartment accompanied by the swelling of the membrane bulge. **D**. The tension dependence on the relative excess membrane area for smaller (1) and bigger (2) compartments. The initial relative excess area was set to $\beta_0 = 0.05$; the size of a small compartment (curve 1) was 0.2 μm; and that of the large compartment (curve 2) was 0.4 μm. Red arrow shows an increase in the PM tension caused by an increase in AC compartment size from 0.2 to 0.4 μm. **E**. The dependence of tension on the size of the compartment base for various compartment's relative excess areas (indicated next to each curve). **D, E**. Membrane bending stiffness and the hydrostatic pressure difference were set to $10^{-19}$ J and 750 Pa, respectively. **F**. Hypothetical pathway from intracellular $Ca^{2+}$ rise to PS-exposure to fusion-promoting deformation of plasma membrane, PM. In the initial state, the ERM (brown squares)-mediated connection between PS (yellow headed lipid)-enriched inner leaflet of PM & underlying actin cortex, AC (green lines) restricts membrane deformations. Transition 1: Lipid scramblase activated by intracellular $Ca^{2+}$ rise releases anionic lipids, including PS, from the inner to outer leaflet of PM, and this results in a local detachment of a fraction of ERM and AC from the PM. Transition 2: Extracellular Anx A5 binds to cell surface PS and further decreases PS content in the inner PM leaflet and ERM- and AC- attachment. Transition 3: Bulging of the PM region disconnected from AC and local increase in tension in this PM region facilitates closer approach between fusing membranes and drives fusion pore formation.

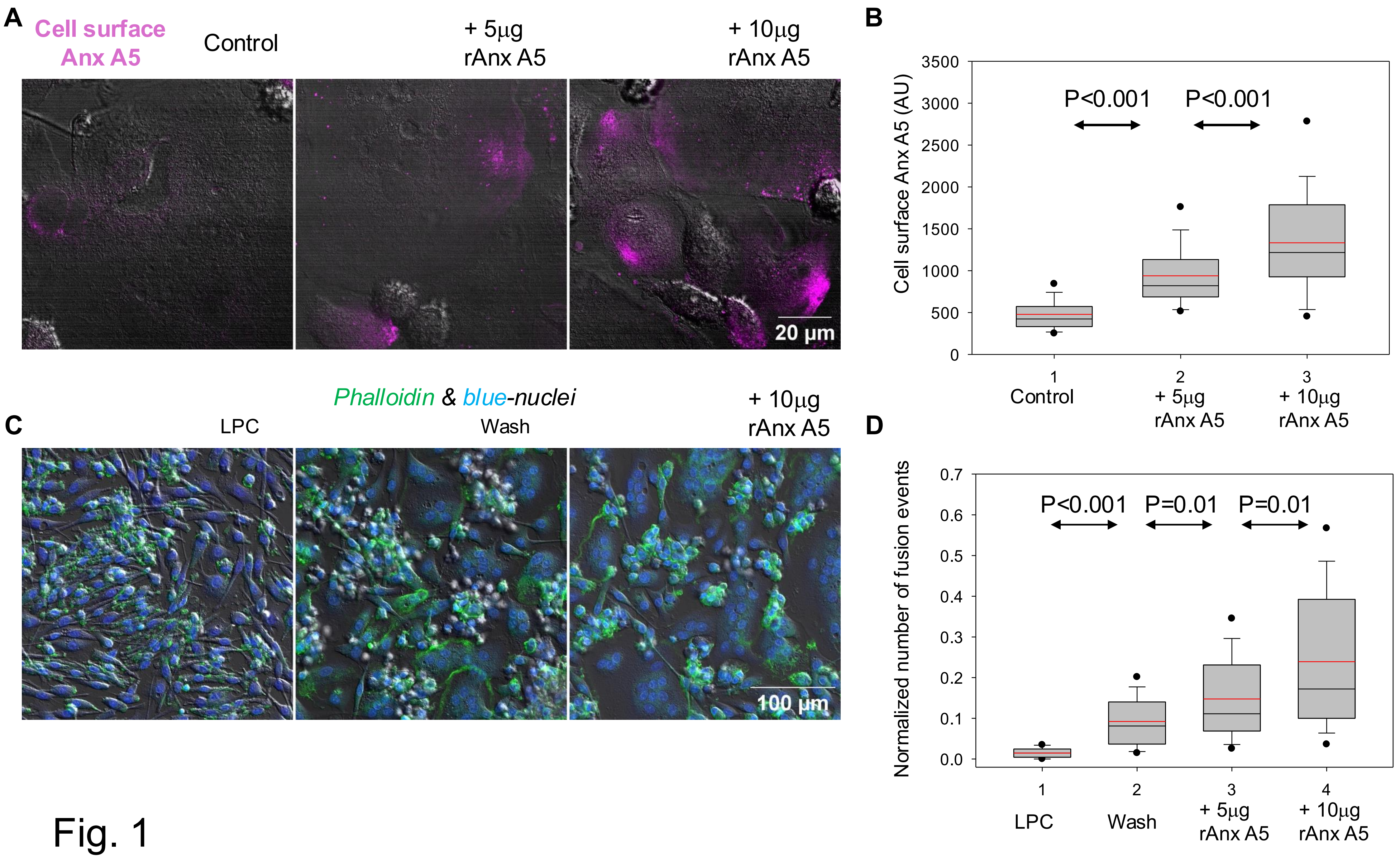
A
Cell surface Anx A5
Control
+ 5µg rAnx A5
+ 10µg rAnx A5
20 µm
B
Cell surface Anx A5 (AU)
3500
3000
2500
2000
1500
1000
500
0
P<0.001
P<0.001
1
Control
2
+ 5µg rAnx A5
3
+ 10µg rAnx A5
C
Phalloidin & blue-nuclei
LPC
Wash
+ 10µg rAnx A5
100 µm
D
Normalized number of fusion events
0.7
0.6
0.5
0.4
0.3
0.2
0.1
0.0
P<0.001
P=0.01
P=0.01
1
LPC
2
Wash
3
+ 5µg rAnx A5
4
+ 10µg rAnx A5

Fig. 1

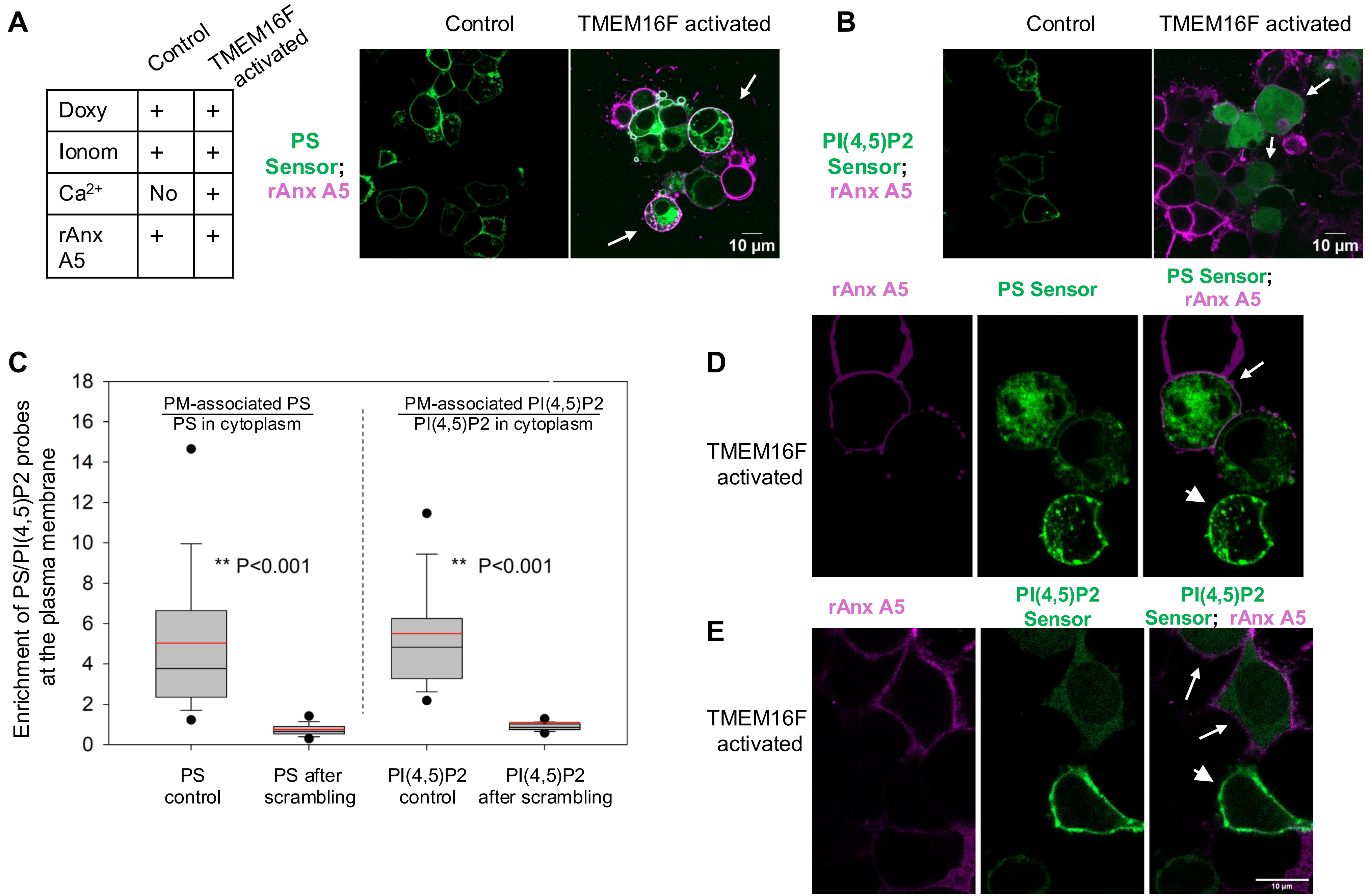
A
Control
TMEM16F activated
Doxy + +
Ionom + +
Ca2+ No +
rAnx A5 + +
Control
TMEM16F activated
PS Sensor; rAnx A5
10 µm
B
Control
TMEM16F activated
PI(4,5)P2 Sensor; rAnx A5
10 µm
C
Enrichment of PS/PI(4,5)P2 probes at the plasma membrane
PM-associated PS / PS in cytoplasm
PM-associated PI(4,5)P2 / PI(4,5)P2 in cytoplasm
** P<0.001
** P<0.001
0
2
4
6
8
10
12
14
16
18
PS control
PS after scrambling
PI(4,5)P2 control
PI(4,5)P2 after scrambling
D
rAnx A5
PS Sensor
PS Sensor; rAnx A5
TMEM16F activated
E
rAnx A5
PI(4,5)P2 Sensor
PI(4,5)P2 Sensor; rAnx A5
TMEM16F activated

Fig. 2

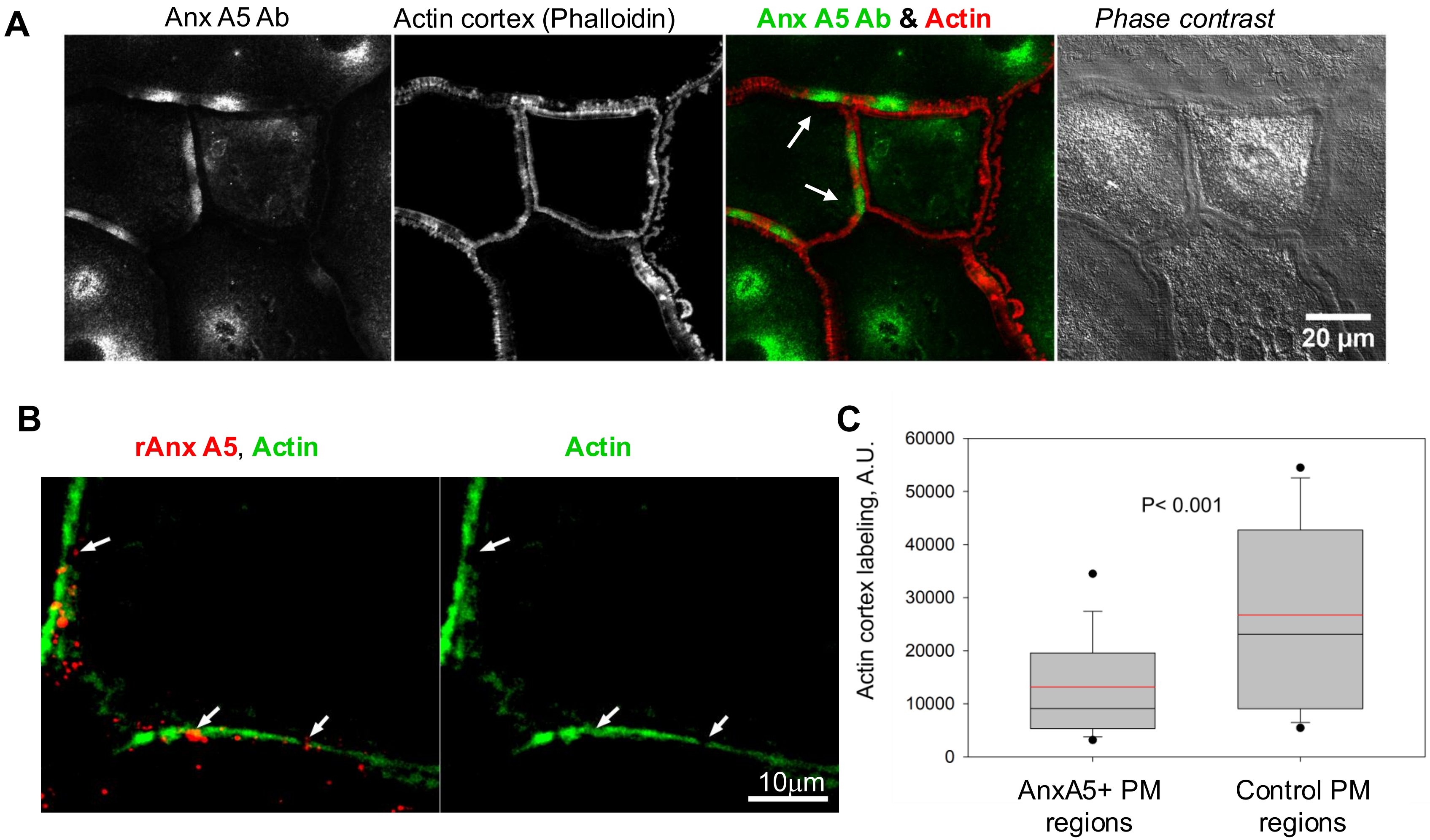
A
Anx A5 Ab
Actin cortex (Phalloidin)
Anx A5 Ab & Actin
Phase contrast
20 µm
B
rAnx A5, Actin
Actin
10µm
C
Actin cortex labeling, A.U.
60000
50000
40000
30000
20000
10000
0
P< 0.001
AnxA5+ PM regions
Control PM regions

Fig. 3

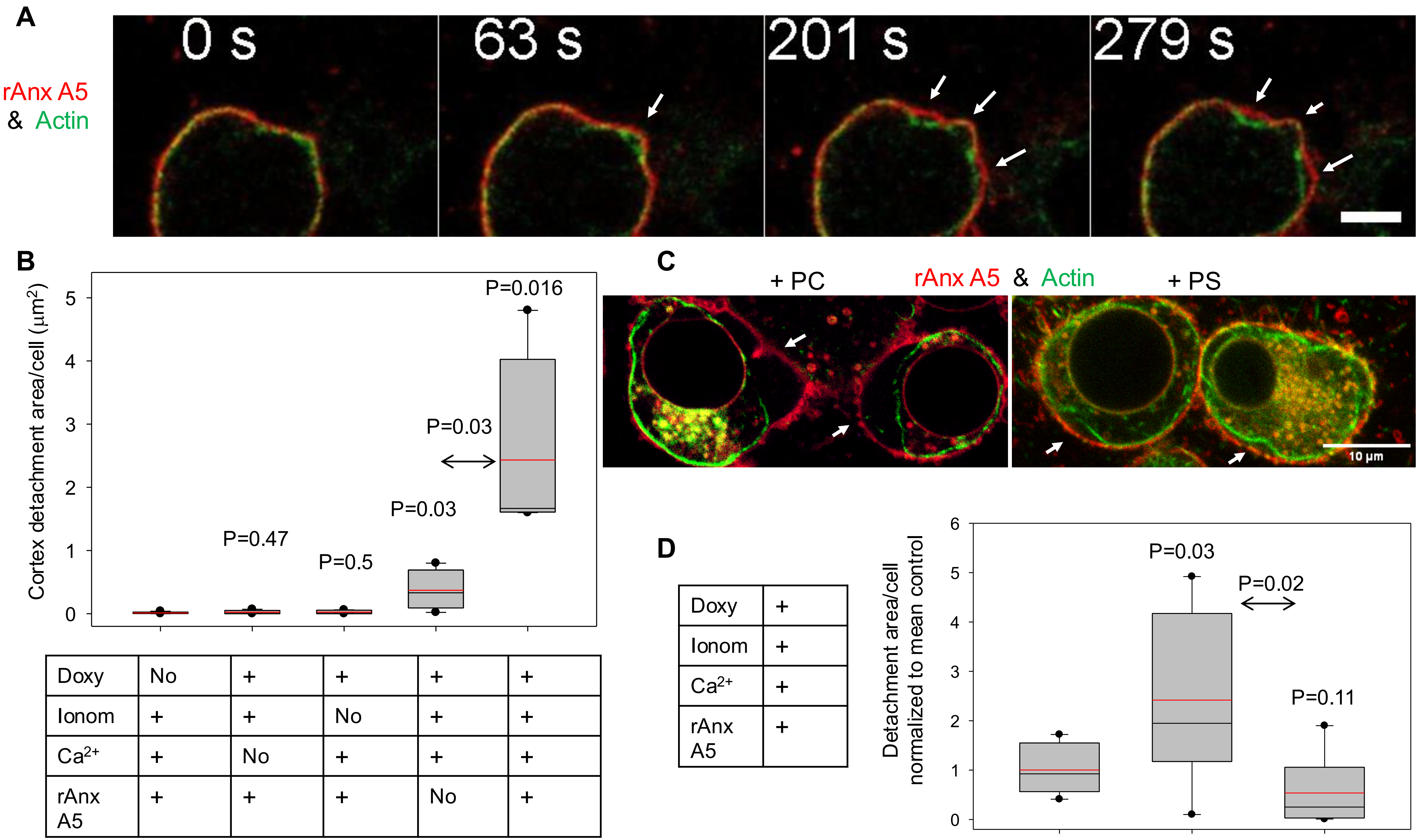


| | | | | | |
|---|---|---|---|---|---|
| Doxy | No | + | + | + | + |
| Ionom | + | + | No | + | + |
| $Ca^{2+}$ | + | No | + | + | + |
| rAnx A5 | + | + | + | No | + |

| | |
|---|---|
| Doxy | + |
| Ionom | + |
| $Ca^{2+}$ | + |
| rAnx A5 | + |

Fig. 4

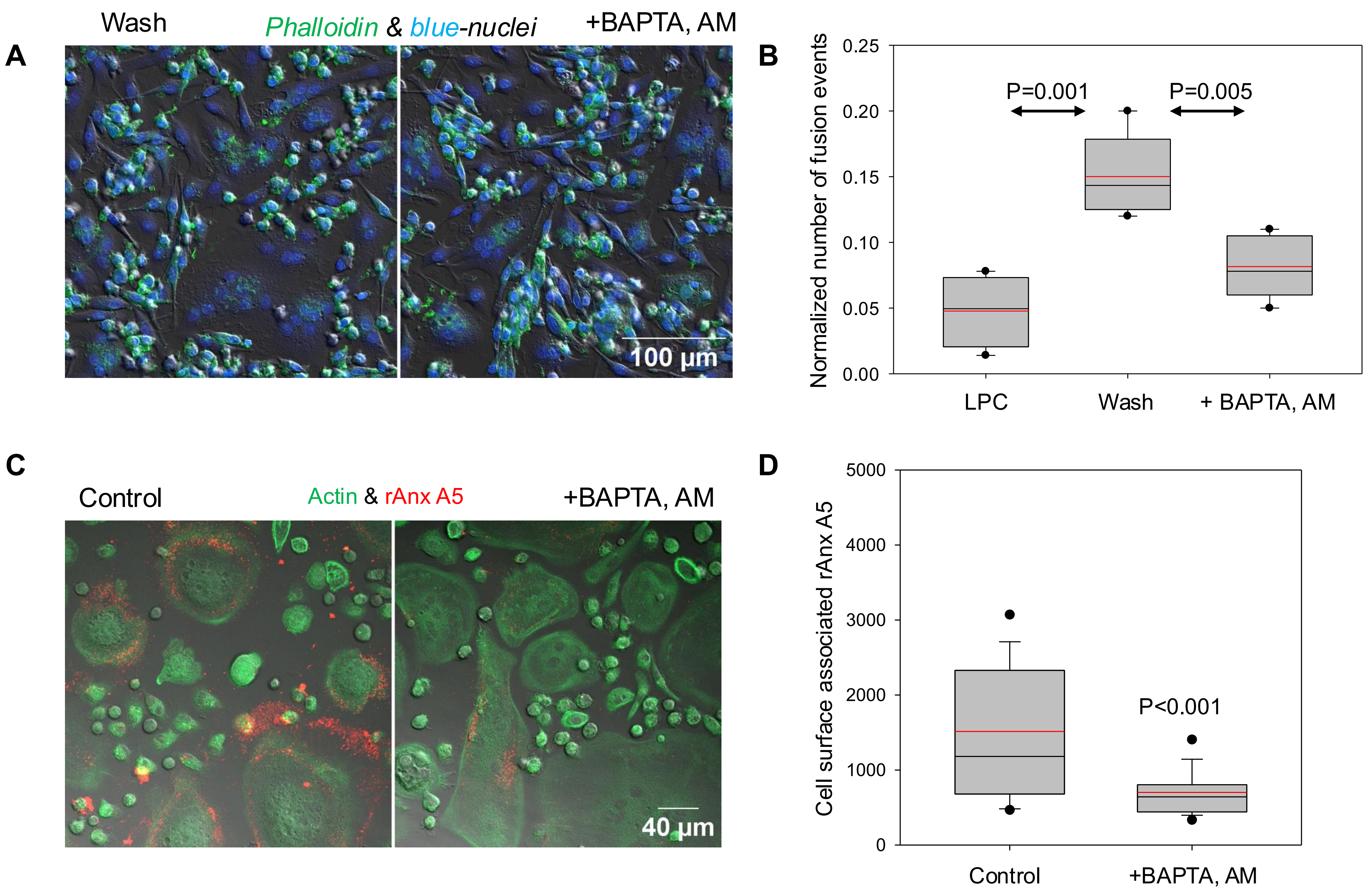
A
Wash
Phalloidin & blue-nuclei
+BAPTA, AM
100 µm
B
Normalized number of fusion events
0.25
0.20
0.15
0.10
0.05
0.00
P=0.001
P=0.005
LPC
Wash
+ BAPTA, AM
C
Control
Actin & rAnx A5
+BAPTA, AM
40 µm
D
Cell surface associated rAnx A5
5000
4000
3000
2000
1000
0
P<0.001
Control
+BAPTA, AM

Fig. 5

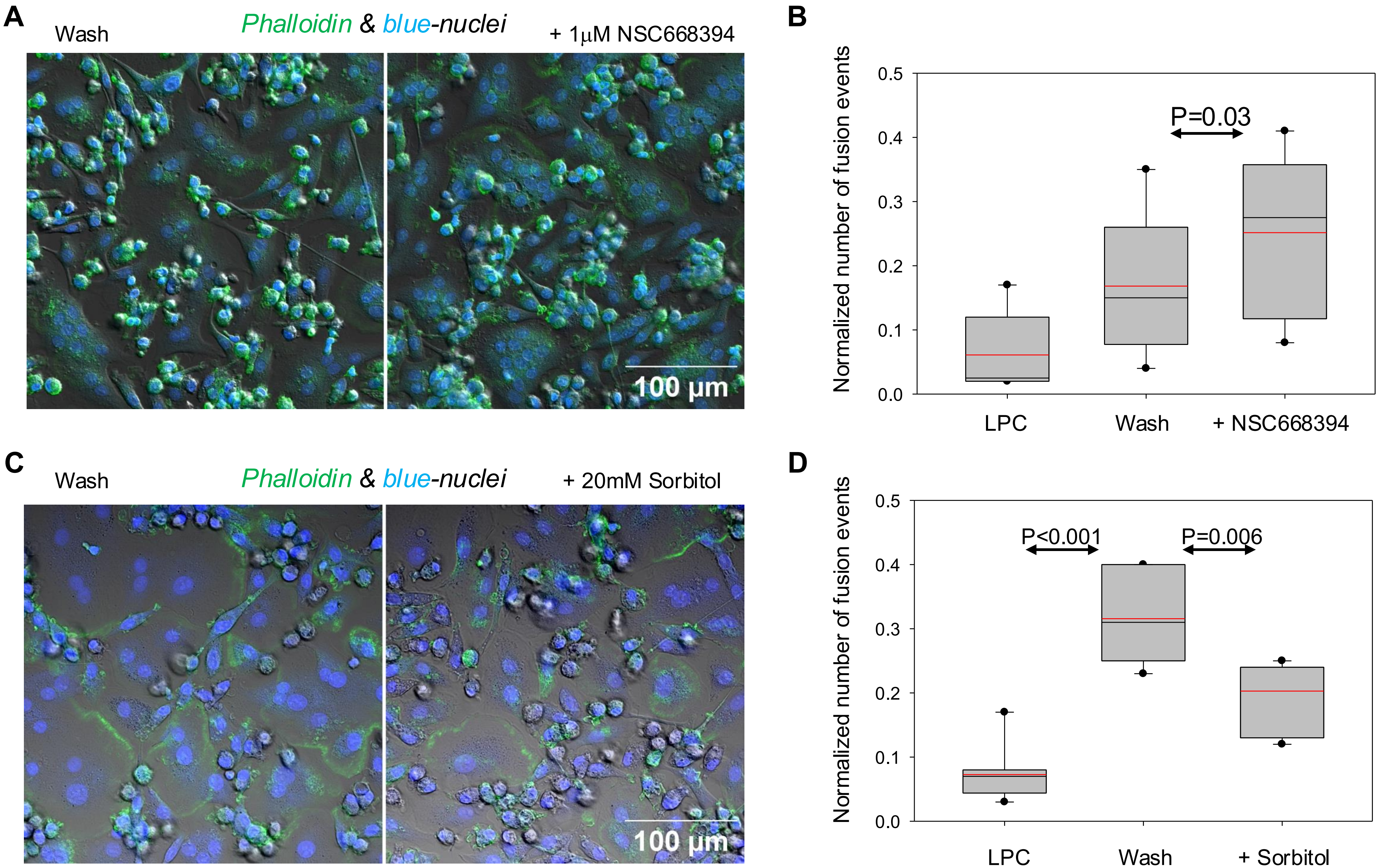
A
Wash
Phalloidin & blue-nuclei
+ 1µM NSC668394
100 µm
B
Normalized number of fusion events
0.0
0.1
0.2
0.3
0.4
0.5
P=0.03
LPC
Wash
+ NSC668394
C
Wash
Phalloidin & blue-nuclei
+ 20mM Sorbitol
100 µm
D
Normalized number of fusion events
0.0
0.1
0.2
0.3
0.4
0.5
P<0.001
P=0.006
LPC
Wash
+ Sorbitol

Fig. 6

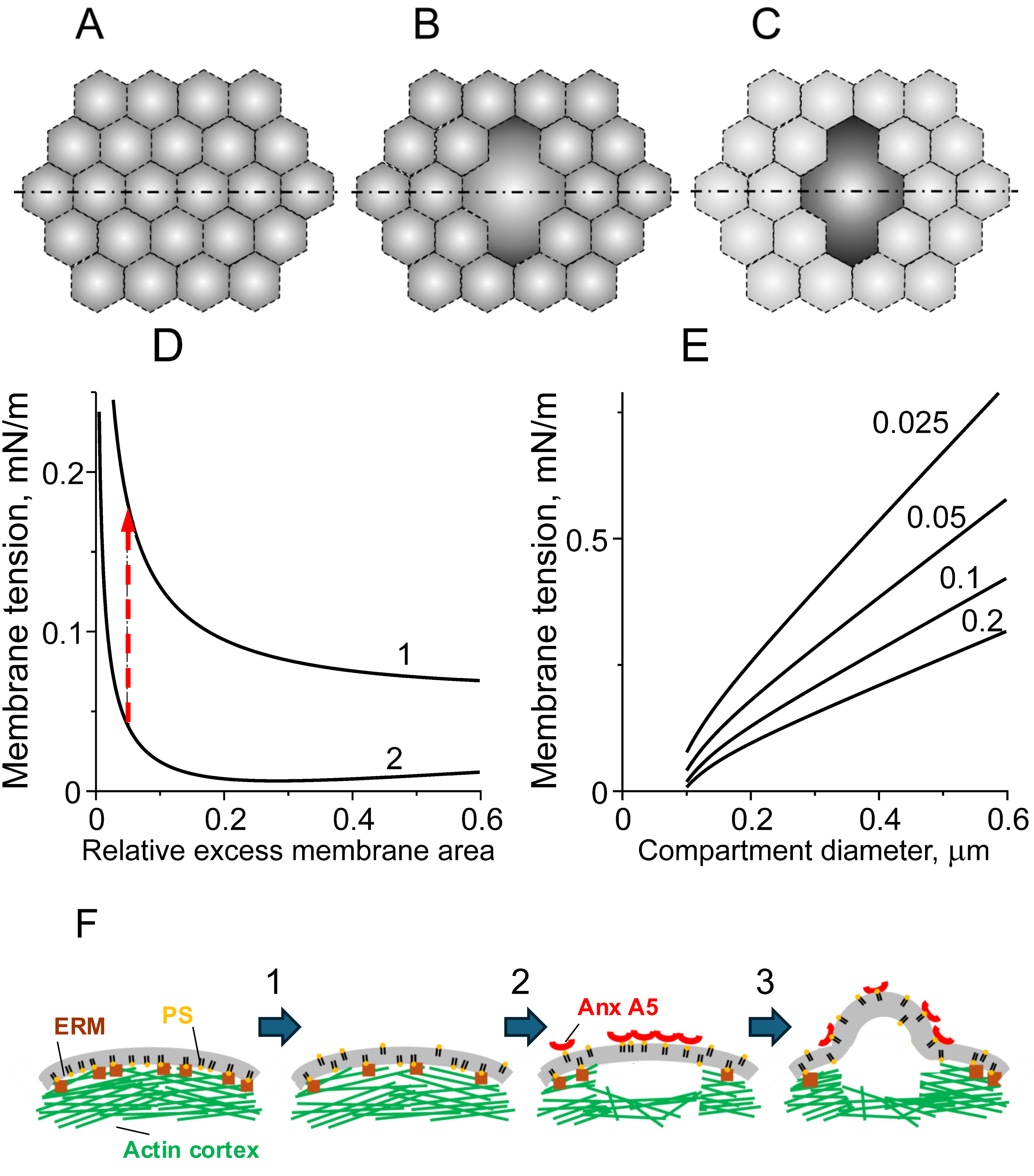
A
B
C
D
E
Membrane tension, mN/m
0.2
0.1
0
1
2
0
0.2
0.4
0.6
Relative excess membrane area
Membrane tension, mN/m
0.5
0
0.025
0.05
0.1
0.2
0
0.2
0.4
0.6
Compartment diameter, µm
F
1
2
3
ERM
PS
Anx A5
Actin cortex

Fig. 7

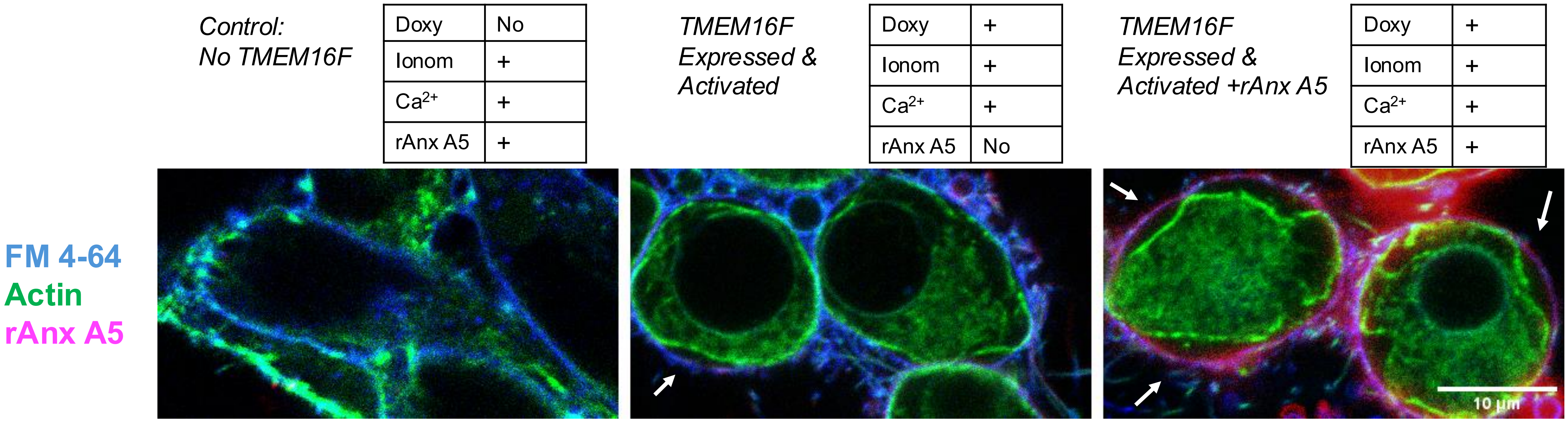


Fig. S1. **Activation of lipid scrambling and rAnx A5 binding promote detachment of actin cortex from plasma membrane.** Representative fluorescence microscopy images of live TMEM16F-HEK cells labeled with Cellmask Green Actin and membrane probe FM4-64 after inducing and activating lipid scramblase TMEM16F in the presence of fluorescent rAnx A5 (magenta) or without it. Prior to activating scramblase, the cells were labeled with F-actin probe (green) and membrane probe FM 4-64 (blue). In control experiments ("No TMEM16F") we did not induce TMEM16F but treated the cells with both Ionomycin or $Ca^{2+}$ application. Arrows mark the actin cortex detachment regions.

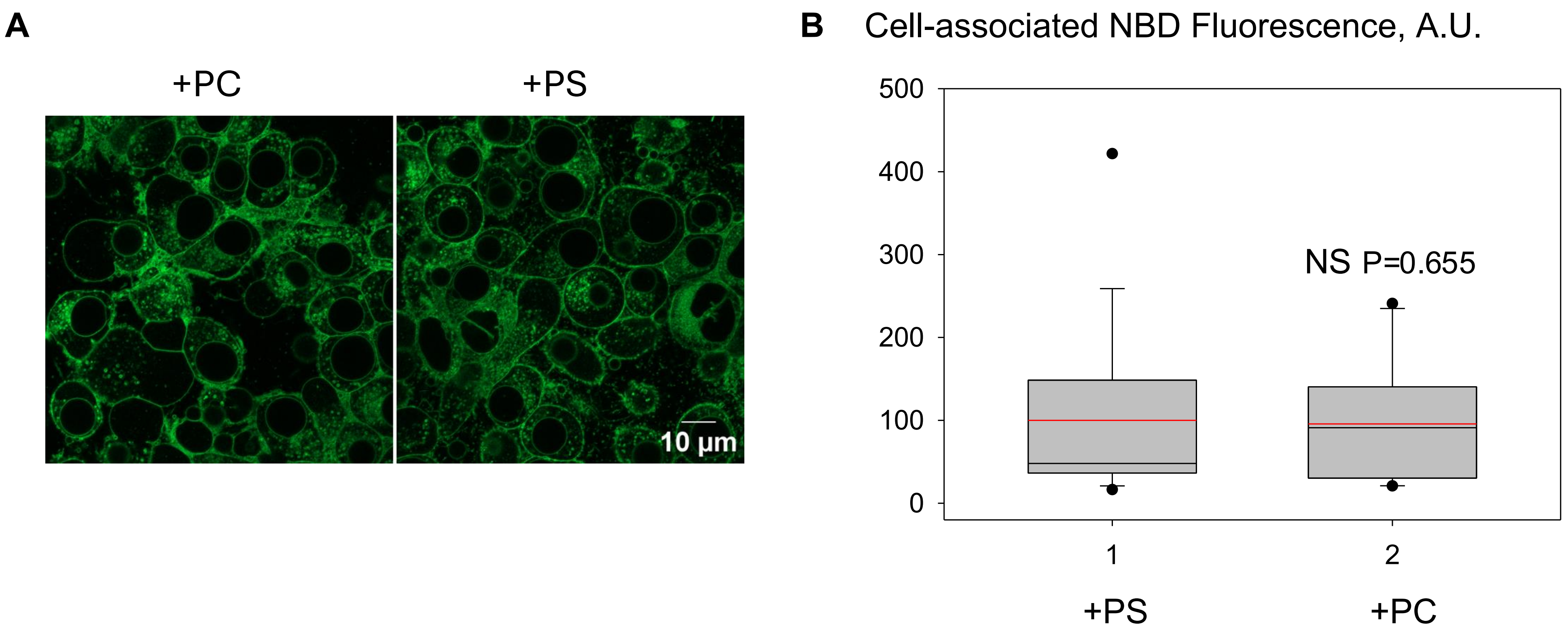


Fig. S2. **Exogenous NBD-tagged phosphatidylcholine (PC) and NBD-tagged phosphatidylserine (PS) associate with the cells in similar amounts.** Cell-associated NBD fluorescence after adding NBD-tagged PC and PS. **A**. Representative images showing cell-associated NBD fluorescence of the TMEM16F-HEK cells with induced and activated scramblase TMEM16F 10 min after application of 10μM exogenous lipids: NBD-tagged PC or NBD-tagged PS, and two washes to remove unbound exogenous lipid. **B**. Quantification of the cell-associated NBD fluorescence in the experiments such as the one in **A**. Data were pooled from 3 independent experiments (at least 10 random imaging fields per condition per experiment and presented as box plots with center lines showing the medians, red lines showing the mean and box limits indicating the 5$^{th}$ and 95$^{th}$ percentiles. Statistical significance was assessed via two-tailed t-test.

### Intracellular $Ca^{2+}$ and rAnx A5

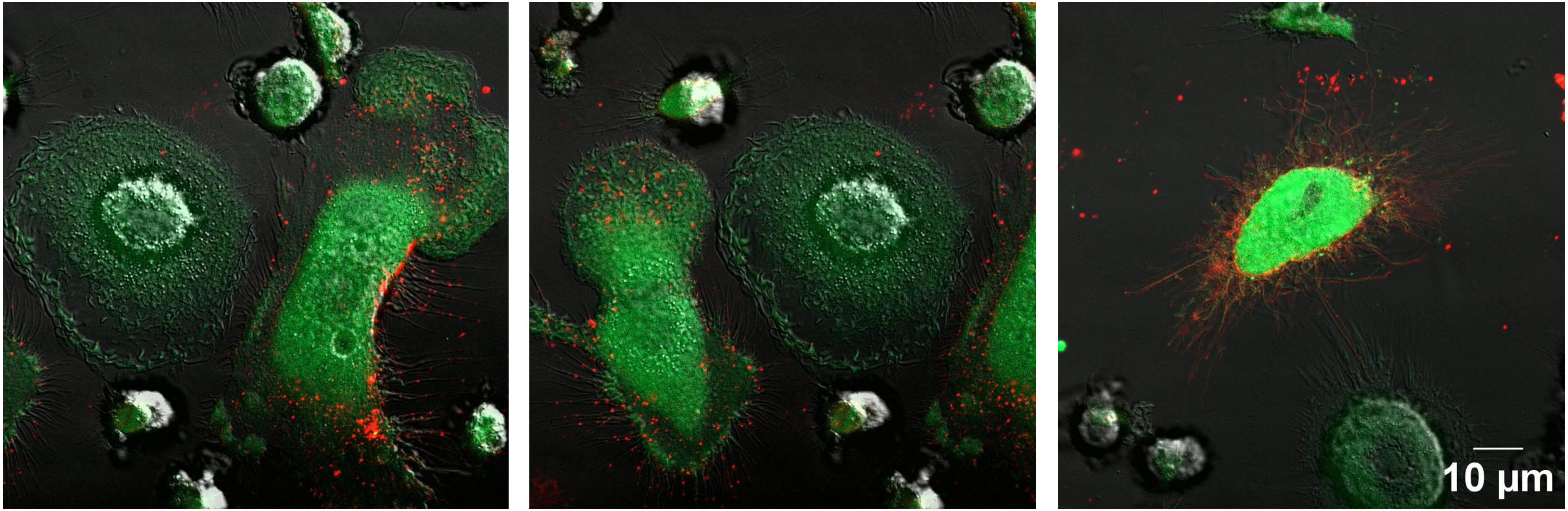


Fig. S3. **Osteoclasts with higher levels of intracellular $Ca^{2+}$ show stronger rAnx A5 staining.** Representative images showing differentiating osteoclasts 4 days post RANKL application. The cells were loaded with 5 µM cell-permeant $Ca^{2+}$ probe Cal-520, AM (green) and treated with fluorescent rAnxA5 (red) to detect intracellular $Ca^{2+}$ and cell-surface PS, respectively.

**Supplementary information**

**Appendix A: Model details**

Here we outline the way of computations of the compartment's tension-area relationships presented in (Figs. 7A-E)

For simplicity, a compartment was approximated by an axisymmetric dome with a circular boundary of radius $a$. The equations of mechanical equilibrium of the compartment's membrane are [1],

$$\frac{d^2J}{ds^2} = -\frac{\cos(\varphi)}{r}\frac{dJ}{ds} + J\gamma_0 + \frac{J}{2}(4K - J^2) = \frac{P}{\kappa}, \quad (1)$$

$$\frac{d\varphi}{ds} = J - \frac{\sin(\varphi)}{r}, \quad (2)$$

where $J$ is the total curvature, $K$ is the Gaussian curvature, $s$ is the distance along the line of membrane cross-section (i.e. the profile line by a plane cut through the axis of symmetry), $\varphi$ is the angle between the tangent to this profile and a horizontal plane, $P$ is the hydrostatic pressure difference between cell interior and extracellular environment, $\kappa$ is the membrane bending stiffness, and $\gamma_0$ is membrane tension in a flat reservoir that would be in equilibrium with the considered membrane compartment.

The total $J$ and Gaussian $K$ membrane curvatures are given by:

$$J = \frac{d\varphi}{ds} + \frac{\sin(\varphi)}{r}, \quad K = \frac{\sin(\varphi)}{r}\left(J - \frac{\sin(\varphi)}{r}\right), \quad (3)$$

respectively.

The boundary conditions at the center of the dome are

$$\varphi(0) = \frac{dJ(0)}{ds} = r(0) = 0. \quad (4)$$

These conditions ensure axial symmetry and a vanishing transverse shear force at the pole of the compartment dome.

At the base of the dome, a zero-angle or zero-torque boundary condition was applied at the dome's boundary, $r = a$:

$$\varphi(s_*)|_{r(s_*)=a} = 0, \quad \text{or} \quad \left.\frac{d\varphi(s_*)}{ds}\right|_{r(s_*)=a} = 0. \quad (5)$$

Here $s_*$ is (a priori unknown) arc length at which the membrane dome radius equals the compartment base radius $a$. The radial coordinate $r(s)$ evolves according to:

$$\frac{dr}{ds} = \cos(\varphi). \quad (6)$$

At a given relative excess membrane area, $\beta$, the value of $\gamma_0$ act as unknown Lagrange multiplier. Alternatively, by solving Eqs. (1), (2) for a given $\gamma_0$ one can compute $\beta$ as:

$$\beta = \frac{2}{\pi a^2}\int_0^{s_*} 2\pi r(s)ds - 1. \quad (7)$$

For small values of $\beta$ and $\varphi$, the non-linear terms in Eq. (1) and (2) can be neglected, allowing an analytical solution using Bessel functions [2]. Alternatively, the equations can be solved numerically using either the shooting method or energy minimization approach such as the finite element-based software Surface Evolver [3] used by Tsaturyan [4].
